# Transmission of ‘*Candidatus* Anaplasma camelii’ to laboratory animals by camel-specific keds, *Hippobosca camelina*

**DOI:** 10.1101/2021.04.02.438174

**Authors:** Joel L. Bargul, Kevin O. Kidambasi, Merid N. Getahun, Jandouwe Villinger, Robert S. Copeland, Jackson M. Muema, Mark Carrington, Daniel K. Masiga

## Abstract

Anaplasmosis, caused by infection with bacteria of the genus *Anaplasma* is an important veterinary and zoonotic disease. The characterization of transmission has concentrated on ticks and little is known about non-tick vectors of livestock anaplasmosis. This study investigated the presence of *Anaplasma* spp. in camels in northern Kenya and whether the hematophagous camel ked, *Hippobosca camelina*, acts as a vector. Camels (*n =* 976) and > 10,000 keds were sampled over a three-year study period and the presence of *Anaplasma* species was determined by PCR-based assays targeting the *Anaplasmataceae* 16S rRNA gene. Camels were infected by *‘Candidatus* Anaplasma camelii*’* occurring from 63 - 78% during the dry (September 2017), wet (June-July 2018), and late wet seasons (July-August 2019). 10 - 29% of camel keds harbored ‘*Ca*. Anaplasma camelii’ acquired from infected camels during blood feeding. We determined whether *Anaplasma* positive camel keds could transmit ‘*Ca*. Anaplasma camelii’ to small laboratory animals via blood-feeding. We show competence in pathogen transmission and subsequent infection in mice and rabbits by both direct detection in blood smears and subsequent molecular identification by PCR. Transmission of ‘*Ca*. Anaplasma camelii’ to mice (8 - 47%) and rabbits (25%) occurred readily after ked bites. Hence, we demonstrate, for the first time, the potential of *H. camelina* as a vector of anaplasmosis. This key finding provides the basis for establishing ked control programmes for improvement of livestock and human health.

**Author summary:** Hematophagous flies such as Tabanids and *Stomoxys*, among other biting flies, are mechanical transmitters of various pathogens such as African trypanosomes and *Anaplasma* species. However, little is known about the role of common camel-specific biting keds (also known as camel flies or louse flies, genus *Hippobosca*) in pathogen transmission. Keds inflict painful bites to access host blood, and in the process may transmit bacterial hemopathogens frequently detected in both camels and their keds. We confirmed by experimental blood-feeding, gene amplification, and amplicon sequencing that camel keds can transmit “*Candidatus* Anaplasma camelii” from naturally-infected camels to healthy mice and rabbits. The high prevalence of camel anaplasmosis throughout the year in northern Kenya could be explained by the infestation camel-specific *H. camelina*, whose capacity as efficient fliers, unlike ticks, promotes disease transmission and maintenance within and among camel herds. Although this study focused on the transmission of *Anaplasma* sp. by camel keds, it is possible that other hemopathogens could also be transmitted by these flies through a similar mechanism. Notably, in the absence of their preferred hosts, keds occasionally bite humans and other vertebrates they come across in order to acquire bloodmeals, and in the process could transmit zoonotic pathogens.

## Introduction

*Anaplasma* species are obligate rickettsial pathogens that proliferate inside red blood cells and cause anaplasmosis in domestic and wild animals. Various *Anaplasma* species such as *Anaplasma marginale, A. centrale, A. platys, A. bovis, A. ovis*, and *A. phagocytophilum* cause huge agricultural losses by severely constraining livestock production and in addition can cause zoonotic infections of humans [1,2]. Clinical signs of anaplasmosis during acute infection include anaemia, pyrexia, reduced milk yield, loss of body condition, abortion, and death [3–5].

*Anaplasma* is transmitted by different species of hard ticks [6] as well as mechanically via contaminated mouthparts of *Stomoxys calcitrans* and Tabanidae, among other biting flies, albeit with reduced efficiency [7,8]. Additionally, pathogen transmission can occur via needles and other veterinary instruments contaminated with fresh infected blood [9]. Mechanical transmission of *Anaplasma* pathogens is thought to be possible only at high parasitaemia [7].

There are increasing reports of camel anaplasmosis with pathogens that include “*Candidatus* Anaplasma camelii” and the related dog pathogen, *Anaplasma platys* [10–12]. Further, *A. platys* continues to be detected in other vertebrate hosts, including humans [13–15], sheep [16], cattle [17], and cats [18]. Close association between humans and their livestock, and co-herding of domestic animals, often in close proximity with wildlife, amplifies chances of spreading vector-borne diseases [15,19]. The veterinary and zoonotic importance of camel anaplasmosis in Kenya, currently with a camel herd of 3.34 million [20], is not well understood.

Camels are kept by nomadic pastoralists in northern Kenya for milk, meat, hides, transport, income from selling milk, and for social capital, for example bride price. Since camels survive readily in harsh arid and semi-arid regions, they are the preferred livestock. However, their productivity is constrained by several factors but chiefly pests and diseases. Other than ticks, keds are the major haematophagous ectoparasitic camel pests and can be found to infest 100% of camel herds throughout the year in northern Kenya. Both male and female keds are obligate blood feeders that stay with their camel hosts, unless disturbed. They mainly infest their vertebrate hosts’ underbelly, although they can be found on the other parts of the body such as neck, ears, hump, and girth [21]. In addition to annoyance and painful bites they inflict while feeding, keds contribute to anaemia, and reduced milk and meat production in camels [22,23]. Keds are members of Hippoboscidae which includes tsetse flies and this family act as vectors of infectious agents such as protozoa [24], bacteria [25], filarial nematodes [26], and viruses [27]. However, *Trypanosoma evansi*-infected camel keds lack competence to transmit trypanosomes to mice [23], and cattle keds, *Hippobosca rufipes*, have been found incompetent to transmit *Anaplasma marginale* from infected cattle to oxen [28].

The main goal of this study was to determine vectorial competence of camel keds for transmission of *Anaplasma* species from naturally infected camels. Our findings show that naturally infected keds are able to transmit ‘*Ca*. Anaplasma camelii’ to laboratory rodents and suggest that keds may act as vectors of wider range of disease agents.

## Materials and methods

### Study area

The present study was carried out in Laisamis (1.6°N 37.81°E, 579 m ASL) located in the south of Marsabit County. Sampling sites were selected along two major seasonal rivers that provide drinking water for livestock, namely: Laisamis River in Laisamis and Koya River located about 29 km south east of Laisamis town (N 01° 23’ 11”; E 37° 57’ 11.7”, 555 m ASL) (Fig 1).

**Fig 1:**
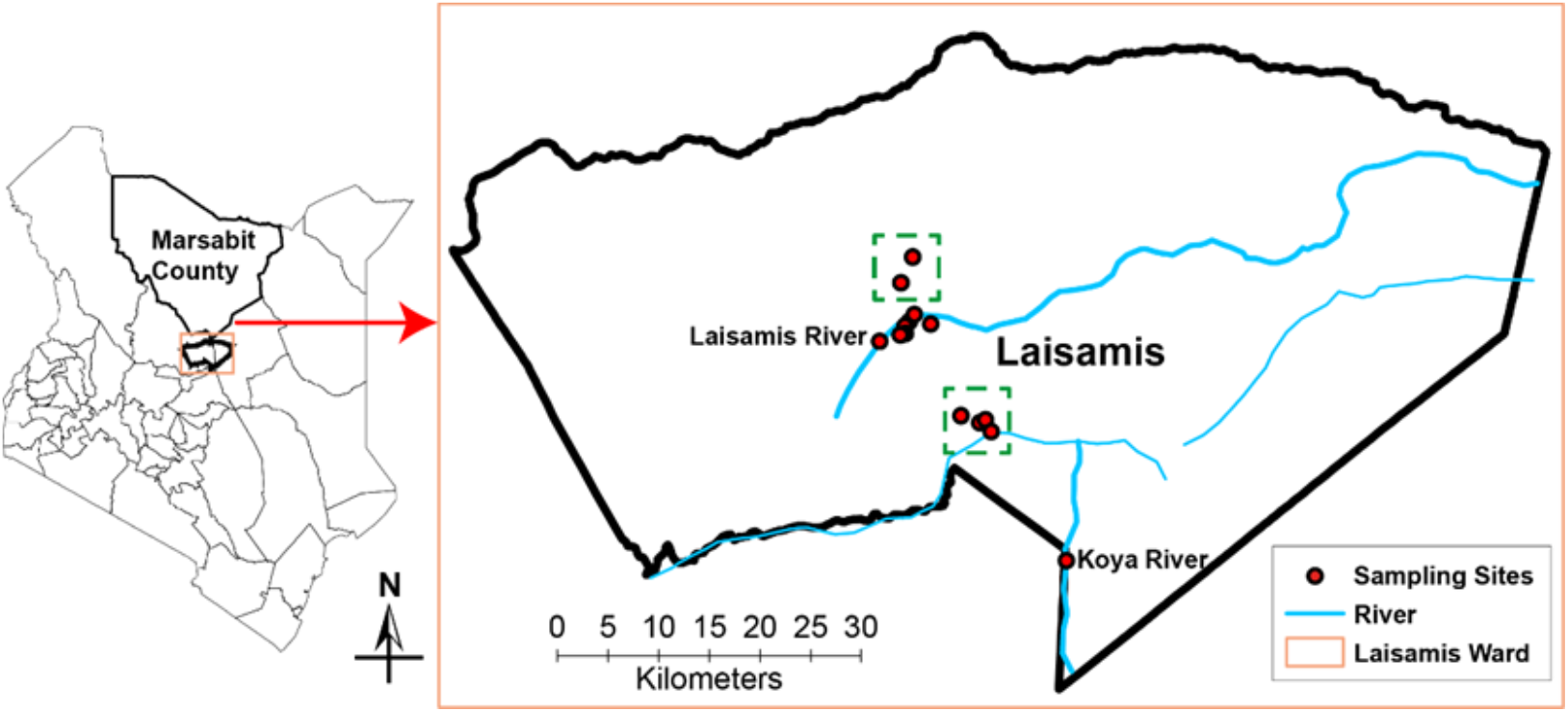
The map of Kenya showing the sampling sites in Laisamis and Koya in Marsabit County.

Marsabit County had a population of approximately 203,320 camels in 2017 with 86% of the household heads, whose major occupation was livestock herding, deriving their livelihoods from the sale of livestock [29].

### Weather conditions

The region is arid and semi-arid and the weather conditions were described in Kidambasi et al [21]. Due to climate change, protracted dry seasons are common resulting in depletion of pastures, decreased livestock productivity, and livestock death [29]. Vegetation dries up soon after wet season except some drought-resistant evergreen trees and shrubs, such as *Acacia tortilis, Cordia sinensis, Salvadora persica, Euphorbia tirucalli*, among others, that camels feed on in dry season.

### Study design and sample collection

This study was cross-sectional in design. Camel blood and ked (*Hippobosca camelina*) samples were randomly collected on daily basis through opportunistic sampling from the available camel herds at the geo-referenced sites. This way, we circumvented lifestyle challenges associated with nomadic pastoralism such as frequent long distance sporadic movement with camels into remote inaccessible areas in search of pastures.

Sampling of blood and keds was initially done during prolonged dry season in September 2017 (*n* = 249 camels). The second sampling to obtain camel blood samples (*n* = 280 camels) was conducted during late wet season in June-July 2018, whereas the third sampling (*n* = 447 camels) was done in July-August 2019 to confirm pathogen transmission by the camel keds and determine establishment of successful infection in test animals.

We lacked information on *Anaplasma* prevalence in Laisamis sub-County for reference in calculating sample sizes due to paucity of historical data, for instance, on the current camel population and production, as well as the burden of pests and diseases, among others. Therefore, we aimed at collecting as many samples as possible during the three main field visits that lasted between 7 – 30 days, and we sampled all camels and keds in randomly selected herds. A herd was defined as a group of camels living together as a unit, often feeding and migrating together. We also surveyed the following in the sampling region (Laisamis); number and sizes of camel herds, and their ownership - mostly pooled together into one herd at family level.

### Ethics statement

We collected camel blood and ked samples, and conducted pathogen transmission experiments in mice with strict adherence to the experimental guidelines and procedures approved by the Institutional Animal Care and Use Committee at the International Centre of Insect Physiology and Ecology. All procedures involving animal handling and use were conducted humanely to minimize pain and distress. CITI training program, on laboratory animals handling and use, was undertaken prior to the approval of the proposed study by *icipe* IACUC (REF: IACUC/ICIPE/003/2018). Sampling during field visits were conducted only after acquirement of informed verbal consent from camel keepers who were unable to read or write, thus written consent was not possible.

### Camel blood collection

The following information was recorded during collection of blood samples; sex, age, count of infesting keds, pregnancy and abortion history, and assessment of the body condition.

Camels were sampled for collection of 5 mL of blood via jugular venipuncture. Blood was drawn into 10 mL EDTA vacutainer tubes (Plymouth PLG, UK) and kept under cold chain at 4°C during collection process. Each labeled sample was transferred into a 2-mL cryotube (Greiner Bio-one, North America, Inc.) and then preserved in liquid nitrogen for transportation to *icipe* laboratories for molecular analysis to detect pathogens.

### Collection of camel keds, *Hippobosca camelina*

We observed that at daytime, collection of keds from camels by hand was difficult because these efficient fliers responded quickly by moving away often landing on the same camel or the next one. In contrast, these flies seemed to be docile at night, thus capturing them off camels was easy. Therefore, whereas blood samples were obtained during the day when camels converged at the watering points along the rivers, keds infesting them were collected later at night between 20:00 – 02:00 hrs. Use of sweep nets for fly collection was discontinued because it frightened camels. Thus, attached keds were hand-picked from camels by 3 – 4 trained field assistants. In order to locate keds on the camel, a spotlight was briefly switched on and then off to minimize light-induced activity (movement) in keds.

Camel keds were strictly collected only from the blood-sampled camel herds in order to allow comparison of the hemopathogens occurring in camels and their keds. It was possible that some keds moved from one camel herd to the next particularly during watering when multiple herds shared a well, even though these flies hardly left their host without disturbance.

Keds were preserved either in the liquid nitrogen or kept at room temperature in absolute ethanol.

### Survey of camel keds; fly populations, and sex

Camel keds were collected from various camel herds and preserved in the absolute ethanol. These collections were sorted out to establish the proportion of flies according to sex. In a separate study to determine the seasonal fly populations, live keds on the camels were counted during dry and wet seasons.

### Morphological identification of camel keds

Keds were morphologically identified at the Zoology Museum of the University of Cambridge (UK) and the Natural History Museum in London using standard morphological keys and by comparison to their preserved ked collections [30].

### Transmission of “*Ca*. Anaplasma camelii” from camels to mice and rabbits by camel keds

The ability of camel keds, *H. camelina*, to transmit *Anaplasma* spp. was studied using mice and rabbits using published protocols with some modifications [23].

Laboratory-reared Swiss white mice (*n* = 21), maintained under clean conditions for 6 - 8 weeks, were used in the first pathogen transmission study conducted in April 2018, while a similar repeat study was done in July 2018 but using caesium irradiated immunosuppressed mice (n = 60; test = 58 and control = 2) to assess the effect of compromised immunity on pathogen acquisition. All other procedures conducted remained the same. In addition, a third independent pathogen transmission study by keds was set up in July-August 2019 using both mice (*n* = 123) and rabbits (*n* = 6). Thus, mice and rabbits were transported to the field sampling sites in northern Kenya to provide bloodmeal for keds freshly collected from camels and in the process finding out any transmitted ked-borne pathogens. Ked feeding schedules were designed as shown in S1-S3 Tables.

Camel keds were collected from a total of 35 camel herds from seven different rural settlements in Laisamis (Fig 1).

### Exposure of camel keds to mice and rabbits for bloodmeal acquisition with concommittant *Anaplasma* transmission

#### Mouse-ked exposure

Freshly collected keds, within 15 min of collection off camels, were directly placed into cages measuring 30(L) × 30(W) × 30(H) cm containing restrained mice to continue blood-feeding on mice for 12 h (*n* = 20 keds/mouse). Ked-exposed mice were allowed to rest for at least 12 h before the next fly exposure to improve chances of pathogen transmission (S1-S3 Tables). All experiments were conducted in a fly-proof environment inside insecticide-free mosquito net in order to exclude other biting flies and also make it easier to recapture keds that escape during handling process. The control mice group was protected from receiving any inadvertent bites from biting flies by wrapping their cages with fly-proof nets.

In order to document evidence of *Anaplasma* infection in cells, pathogen transmission study was replicated for the third time under same conditions as described before, with minor changes on the host-ked exposure, but this time using both mice and rabbits as bloodmeal sources (S3 Table). Slight modification was adopted to improve feeding success in flies by placing fly cages, usually covered on both sides by a netting, on the shaved body parts of animals that are restrained by hand during fly feeding, rather than introducing wire-mesh restrained mice into the fly cages.

#### Rabbit-ked exposure

Restrained test rabbits (n=4) were exposed to ked bites to feed repeatedly on each rabbit with a 2-day interval in between successive feeds (n=40 keds/rabbit) in order to increase chances of pathogen transmission, if at all possible. Ked inflicted bites on the rabbit ears and on shaved region on the back. The control group (n=2) was protected from biting flies throughout the study. All mice and rabbits were screened regularly for *Anaplasma* infection following ked bites.

### Screening of mice and rabbits post-ked bites

Blood samples were collected at intervals for pathogen screening in both control and test mice and rabbits. Towards the end of the follow up pathogen detection studies, all mice were anaesthetized prior to collection of about 1mL of blood from the heart using 1 mL syringes containing 200 µL of Carter’s Balanced Salt Solution with 10,000 Units/L heparin sodium salt.

Blood samples were also obtained from rabbits by pricking the ear vein with a lancet followed by sample collection in heparinized capillary tubes.

Wet blood smears were prepared for staining to detect *Anaplasma* infection and total DNA was extracted from blood samples for time-course detection of *Anaplasmataceae* by PCR-based assays as described below.

### Field’s staining of blood smears for parasitological examination for detection of *Anaplasma* spp

Rapid Field’s staining of thin blood films was done to detect *Anaplasma* species in the mice (n = 8 smears), and rabbit blood smears (n = 8 smears), as well as fixed camel samples (n = 13 smears) prepared during field sampling.

The tip of mouse-tail and rabbit ear was sterilized with 70% v/v ethanol, and a vein was pricked using sterile lancet to collect a drop of blood on a microscope slide for preparation of thin blood smears. Each lancet was used only once to avoid iatrogenic transmission of *Anaplasma* in test mice and rabbits.

During field collection of samples, a drop of blood from each camel was placed on to a slide for preparation of thin blood smears. Air – dried thin blood smears were labeled, air-dried, fixed for 1 minute in methanol, and then air-dry the film prior to staining. Field’s staining was performed by flooding the slide with 1 mL of Field’s stain B (diluted 1 in 4 with distilled water; stain B contains methylene blue and Azure dissolved in a phosphate buffer solution) followed immediately by addition of equal volume of Field stain A (undiluted; stain A has Eosin Y in a buffer solution) using Pasteur pipette. Field’s stains A and B were mixed well using Pasteur pipette and the staining reaction was allowed for 1 minute, followed by rinsing of slides in distilled water. The slides were air-dried for imaging to morphologically identify intracellular species of *Anaplasma* pathogens using digital camera ZEN lite 2012 mounted on to a compound microscope equipped with 60× and 100× oil-immersion objectives [31]. Ten (10) microscopic fields were counted for each slide and average percentage bacteremia levels determined as in formula below;

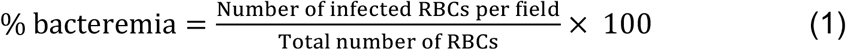

### Molecular identification of *Anaplasma* species

#### DNA extraction from blood and ked samples

Each camel ked, with an average body weight of 100 mg, was surface-sterilized with 70% ethanol and briefly allowed to air-dry on a paper towel. Each whole ked was then placed into a clean 1.5-ml microfuge tube containing sterile 250 mg of zirconia beads with 2.0 mm diameter (Stratech, UK) and ground in liquid nitrogen in a Mini-Beadbeater-16 (BioSpec, Bartlesville, OK, USA) for 2 min. Total genomic DNA was extracted from each ked lysate and blood (camel, mouse, and rabbit) using DNeasy Blood & Tissue Kit (Qiagen, Hilden, Germany) following the manufacturer’s instructions.

### PCR - HRM, purification of PCR amplicons and gene sequencing

Detection of *Anaplasma* spp. was done using conventional PCR followed by high resolution melting (PCR - HRM) analysis in a HRM capable RotorGene Q thermocycler (Qiagen, Hannover, Germany). Screening for *Anaplasma* spp. by PCR - HRM was done using *Anaplasma*JV-F and *Anaplasma*JV-R primers (Table 1) as previously described by Mwamuye *et al*., 2017.

**Table 1:**
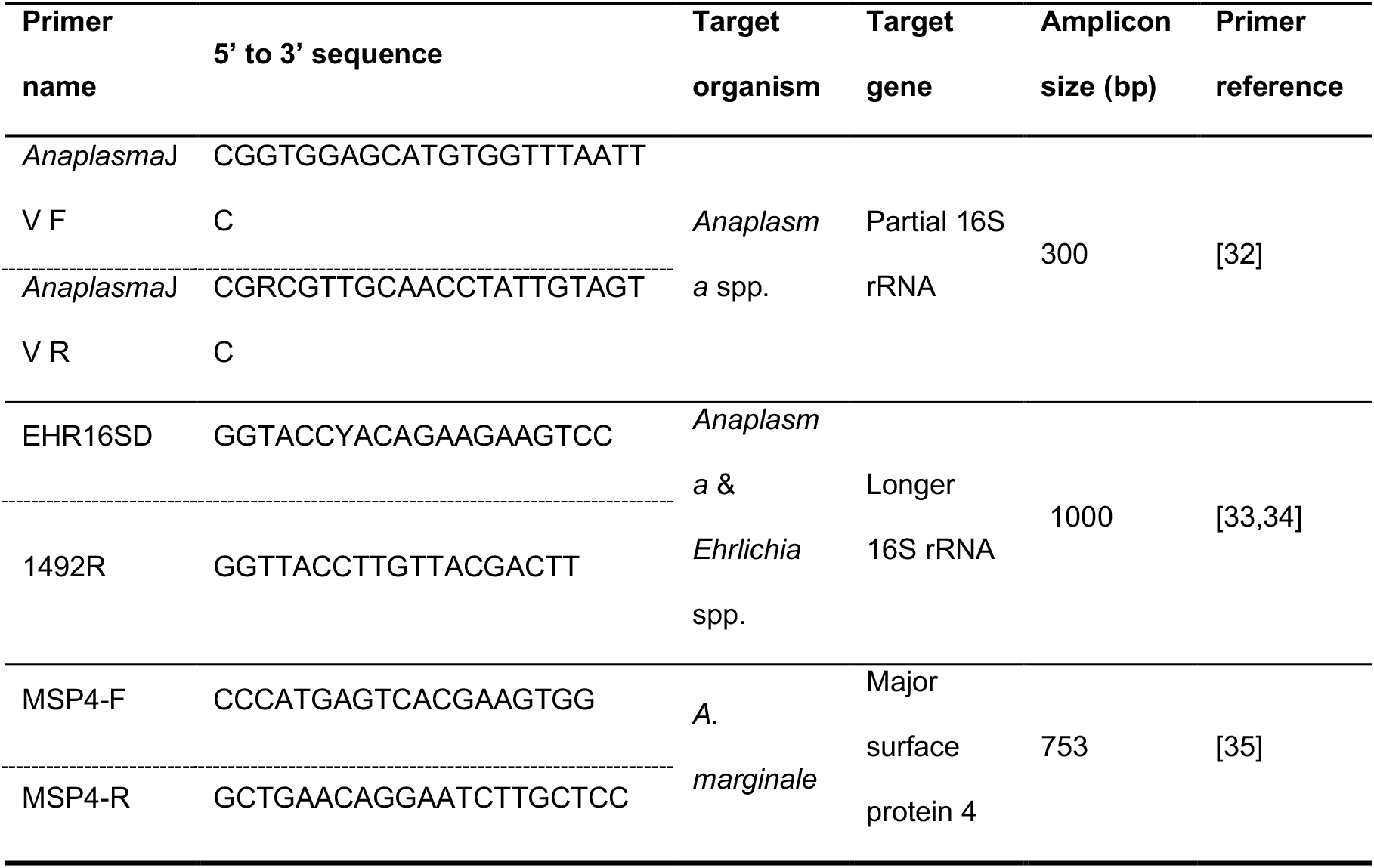
Primers for PCR amplification.

The PCR-HRM assays were carried out in 10 μL reaction volume, containing 2.0 μL of 5× HOT FIREPol EvaGreen HRM mix (no ROX) (Solis BioDyne, Estonia), 0.5 μL of 10 pmol of each primer, 6.0 μL PCR water and 1.0 μL of template DNA. The amplification conditions included an initial enzyme activation at 95°C for 15 min, followed by 10 cycles at 94°C for 20 sec, step-down annealing for 25 sec from 63.5°C to 53.5°C decreasing the temperature by 1°C after each cycle and an extension step at 72°C for 30 sec; followed 25 cycles at 94°C for 25 sec, annealing at 50.5°C for 20 sec, extension at 72°C for 30 sec, and a final elongation at 72°C for 7 min. Immediately after PCR amplifications, HRM curves of the amplicons were obtained by increasing temperature gradually from 75°C to 95°C at 0.1°C/2 sec increments between successive acquisitions of fluorescence. Negative and positive controls were included in all PCR - HRM assays. The HRM profiles were assessed using Rotor-Gene Q Series Software 2.3.1 (Build 49). Changes in fluorescence with time (dF/dT) were plotted against corresponding changes in temperature (°C). PCR-HRM amplified only smaller region of 16S rRNA gene, therefore, representative samples with unique peaks were further PCR-amplified by conventional PCR for sequencing longer fragment of the *Anaplasma* 16S rRNA gene of ∼1000-bp using EHR16SD and 1492R primers [33,34] (Table 1). Longer sequences enabled species identification and were used in phylogenetic analysis. The PCRs were performed in 15 μL reaction volumes that included 5.0 μL PCR water, 1.0 μL of DNA template, 3 μL of 5 x HOT FIREPol® Blend Master Mix (Solis Biodyne, Estonia) and 0.5 μL of 10 μM EHR16SD and 1492R primers (Table 1). The cycling conditions consisted of: 95°C for 15 min; 2 cycles of 95°C for 20 sec, 58°C for 40 sec, and 72°C for 90 sec; 3 cycles of 95°C for 20 sec, 57°C for 30 sec, 35 cycles of 95°C for 20 sec, 56°C for 40 sec and 72°C for 80 sec, and a final extension at 72°C for 10 min. The amplification was carried out in a ProFlex PCR system (Applied Biosystems by life technologies).

PCR amplicons were resolved on 1% ethidium bromide-stained agarose gel at 80 V for 1 hr. The DNA was visualized under UV transillumination and the target bands excised and purified from the gel using QIAquick PCR purification kit (Qiagen, Valencia, CA) following manufacturer’s instructions and sequenced by Sanger method (Macrogen Europe, Amsterdam). The identity of *Anaplasma* was confirmed to be ‘*Ca*. Anaplasma camelii’ and no cases of mixed *Anaplasma* infection were found. The sequences generated were deposited in the GenBank with accession numbers as follows; (i) Camel keds: MK754149-MK754151 and MT510535-MT510537, (ii) Camels: MK754152-MK754154, MT510527, MT510529, MT510531, MT510532, and MT510534, (iii) Mice: MK754155-MK754160 and MT510538, and (iv) Rabbit: MT510539. The GenBank accessions for the longer ∼1000-bp sequences used for phylogenetic analysis include; MK388294-MK388300, MT510528, MT510530 and MT510533 (sequenced in camel) & MK388301 (sequenced in test mouse after ked feeding bites).

### Data analysis

The geo-referenced data on sampling locations was fed into a GIS package (ArcGIS v 10.6) and the maps were generated. Data on ked catches was entered into Microsoft Excel spreadsheet, version 12.3.1 and exported to R software, version 3.5.1 for analysis.

Keds were collected longitudinally for a 3-year period spanning wet, late wet, and dry seasons. To determine whether the observed keds infestation on camels (response variable), and the likelihood of pathogen transmissions was influenced by both environmental (seasonal conditions; dry, wet, late wet) and host-specific factors (age; young or, mature), or sex (male, female), we fitted a generalized linear model (*glm*) assuming a Poisson error distribution with *log* link function. The influence of the following covariates was further analysed; exposure frequency and ked average numbers on pathogen transmission status (positive = 1, negative = 0) of experimental mice. Differences between various variables were compared using analysis of deviance F statistics with Chi-square test. For all the analyses, a *p* value of less 5% (*p* < 0.05) was considered significant. ANOVA was used to determine variations in keds infestation on various livestock species. All study nucleotide sequences were edited in Geneious Prime 2019.1.1 software version (created by Biomatters) using the MAFFT plugin [36] and aligned with related sequences identified by querying in the GenBank nr database using the Basic Local Alignment Search Tool (www.ncbi.nlm.nih.gov/BLAST/).

Phylogenetic and molecular evolutionary analyses were performed using MEGA5 software [37]. Phylogenetic trees were constructed using Neighbor-Joining method with 1000 bootstrap replicates. *Wolbachia* endosymbiont (family *Anaplasmataceae*) was included in the phylogenetic tree as outgroup.

## Results

### Survey of camel keds, *Hippobosca camelina* in northern Kenya

#### (a) Keds infestation on different herds

Identification of keds collected from camels, sheep, goats, cattle, dogs, and donkeys showed that *H. camelina* was only found on camels and not on other domestic animals, suggesting that *H. camelina* (Fig 2) is camel-specific.

**Figure 2:**
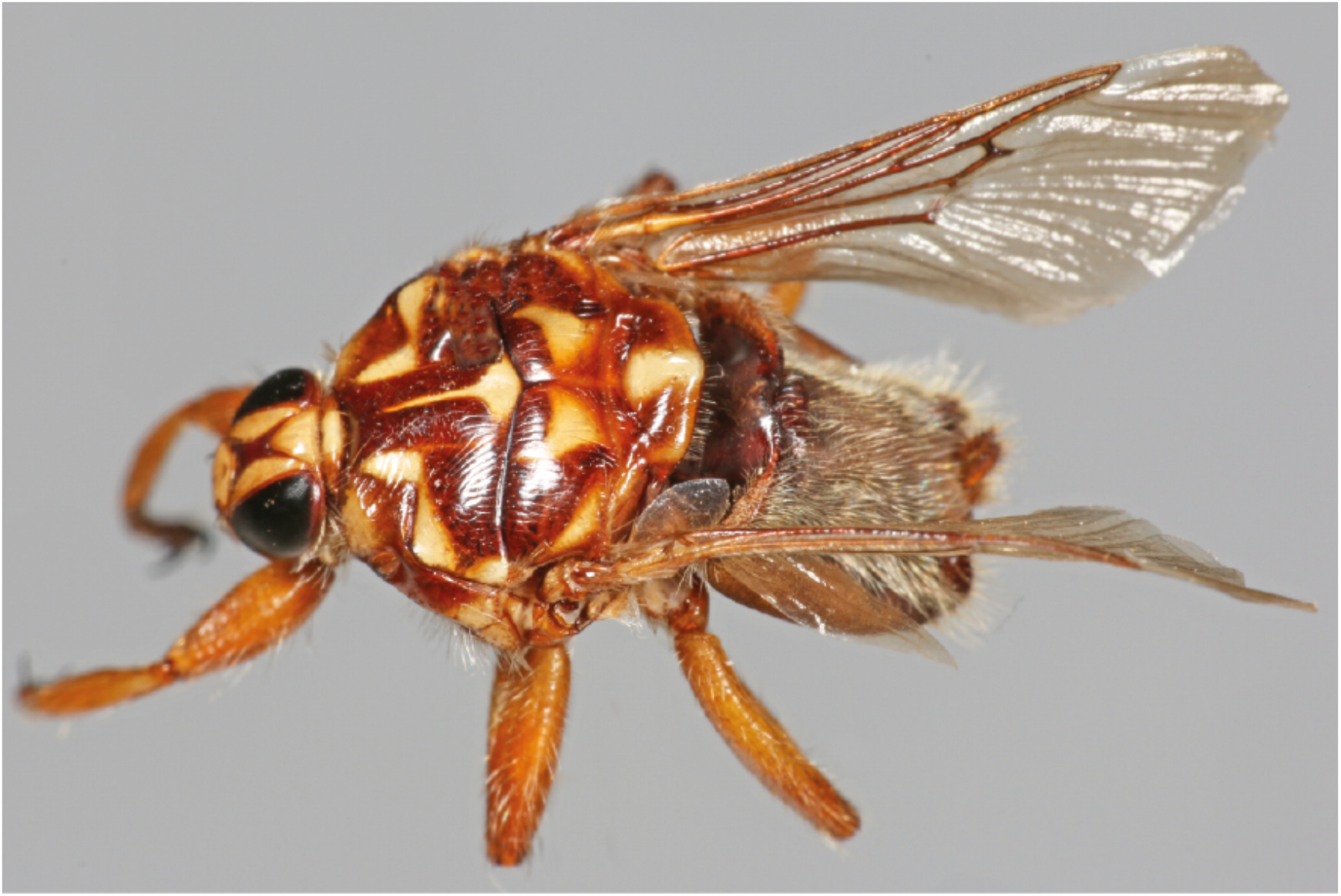
Hippobosca camelina.

The highest mean ked numbers were found on camels with fewer on sheep and goats, significantly varying across the sampled animals (ANOVA, F(5, 273) = 12.64, p < 0.0001). The mean counts of keds on camels were relatively higher compared to those collected from cattle (Welch two-sample t-test; t = −7.2872, df = 206, p < 0.001), but not donkeys (t = 1.884, df = 16, p = 0.078). Similarly, more keds were collected from goats than sheep (t = 2.3442, df = 47, p = 0.0234). Moreover, we collected more keds on cattle in comparison to sheep (t = −5.4735, df = 58, p < 0.001). No significant variation was found between mean ked counts on donkeys and dogs (t = 0.8346, df = 15, p = 0.4174), and between camels and dogs (t = 0.4602, df = 11, p = 0.6545) (Fig 3A).

**Fig 3:**
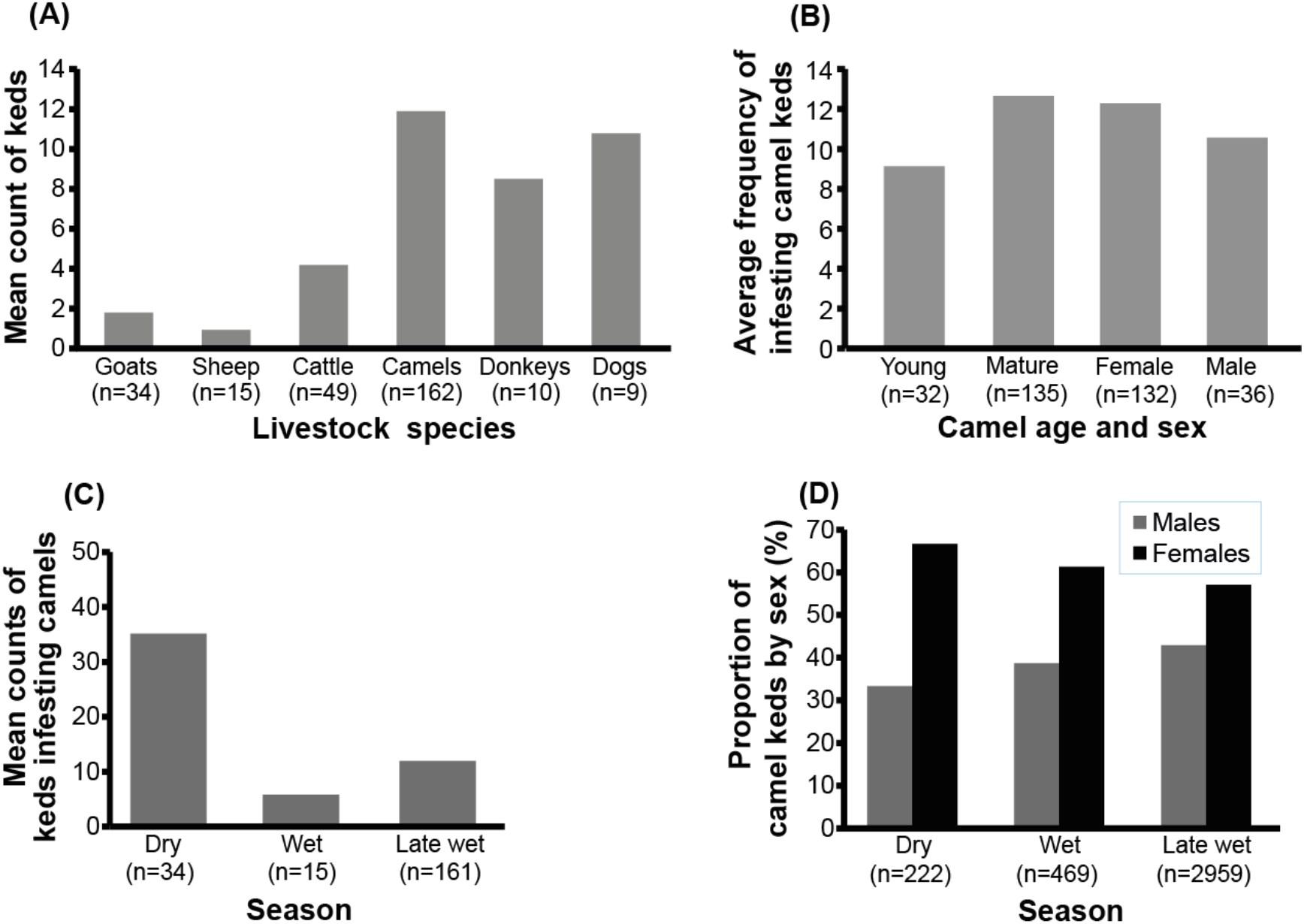
Baseline survey data of livestock keds in northern Kenya.

**(A) Ked infestation in domestic animals**. Keds of other domestic animals, including goat, sheep, cattle, donkey, and dogs, were also sampled alongside camels to compare fly densities in various livestock species and determine whether keds show host-specific preferences. *H. camelina* was only collected on camels, but not from other livestock showing its camel-specific preference. Highest mean ked infestation was recorded in camels, followed by dogs, and then donkeys.

**(B) The influence of host age and sex on the preference of keds to infest camels**. Mature camels had higher average numbers of keds as compared to the young camels.

**(C) Shows comparison of the proportion of male and female keds**. The keds were randomly collected from camels during three seasons: dry, wet, and late wet seasons.

**(D) Seasonal variation of ked populations**. The proportion of female keds was higher than the male flies sampled across the three seasons.

We observed that keds were difficult to collect during the daytime as they quickly moved away, unlike at night when they seemed docile in the absence of light. *Hippobosca camelina* was present: (i) in all of the camel herds suveyed, (ii) throughout the year, and (iii) nearly all camels in herds were infested with keds.

#### (b) Keds infestation on camels of different ages

Ked infestation significantly varied with the camel age; young camels under two years of age were infested by 1 - 35 keds/camel with an average fly number of 9, whereas camels older than 2 years were infested by 0 - 82 keds/camel with mean of 13 (glm; F_(1,528)_ = 4375, *p* < 0.001). In contrast, the number of camel keds did not differ significantly with the camel sex (glm; F_(1,527)_ = 4374, *p* = 0.5508) (Fig 3B).

#### (c) Seasonal variations of keds infestation on camels

We sampled camel keds in the field to determine the average fly numbers per camel during the dry, wet, and late-wet seasons. The average number of keds varied significantly with seasons (glm; F_(1,699)_ = 7835, *p* < 0.001). The highest number of keds were present in the middle of dry season, reaching up to 100 keds/camel. In contrast, during the wet season, the number of keds/camel reduced to an average of 7 keds/camel (*n* = 14 camel herds). Towards the late-wet season in June-July 2018, ked numbers increased gradually to an average of 10 keds/camel. This rising trend in the number camel keds continued until reaching peak numbers of 80 - 100 keds/camel in the dry season (Fig 3C). This seasonal variation in the mean number of keds on the camel host repeats every year.

#### (d) Proportion of male and female keds in dry, wet, and late wet seasons

We found more female than male keds irrespective of the season of the year in all randomly collected keds from various camel herds in different geographical locations sampled at different times. Overall, the ked collections comprised of twice as many female flies than males; i.e. 66.7% females, and 33.3% males in dry season (*n* = 222 keds) (Fig 3D). This 2:1 ratio of female:male keds was consistent in all seasons (ANOVA, F_(1,4)_ = 0.1024, *p* = 0.765), (wet season, t = 4.4231, df = 1, *p* = 0.1416; dry season, t = 3, *p* = 0.2048; late-wet season, t = 7.0285, df = 1, *p* = 0.08997).

### Molecular detection of *Anaplasma* spp. in camels and their keds

#### (a) *Anaplasma* in camels

DNA was extracted from camel whole blood to test for the presence of *Anaplasma* using PCR amplification of 16S rRNA gene marker. The prevalence of *Anaplasma* in camels was generally high across all the sampling seasons. The rate of infection during dry, wet, and late wet seasons was 70.3% (*n* = 175/249), 63.9% (*n* = 179/280), and 77.9% (*n* = 348/447), respectively (S4 Table). We found ‘*Ca*. Anaplasma camelii’ sequences from study camels share 100% identity with reference sequences (GenBank accessions: KF843824 and KX765882) sequenced from camels in Saudi Arabia and Iran, respectively.

#### (b) *Anaplasma* in camel keds

Keds were collected from the same herds (collection at level of individual camels was not possible as keds often move to a different host when disturbed) and DNA prepared from individual fly to detect presence of *Anaplasma* by PCR amplification of 16 rRNA gene marker. The prevalence of “*Ca*. Anaplasma camelii” in keds varied with season, with the percentage prevalence being: dry (9.9%), wet (20.8%), and late wet (28.9%) (S4 Table).

### Experimental transmission of *Anaplasma* from camels to laboratory animals

Mice and rabbits were repeatedly exposed to ked bites freshly collected from camels. Field’s stained thin film blood smears were used to detect the presence of *Anaplasma* in red blood cells of the experimental laboratory animals. This approach readily detected *Anaplasma* in camels (Fig 4A) and was also successful when used with test mice (Fig 4B), and rabbits (Fig 4C). The percentage red blood cells containing *Anaplasma* was 1.6% in camels, 2.7% in mice, and 3.0% in rabbits, respectively (Fig 4).

**Fig 4:**
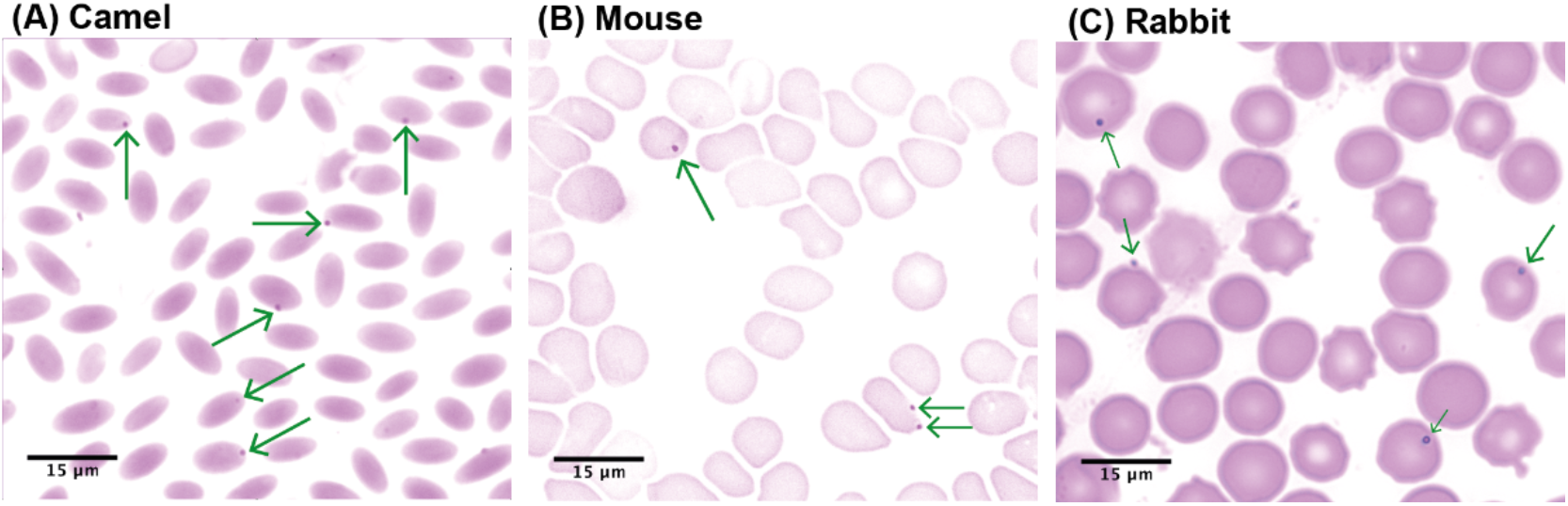
Field’s staining showing *Anaplasma* sp. infections in representative thin film blood smears. **(A)** Naturally infected dromedary camel, **(B)** Experimental mouse exposed to ked bites, **(C)** Experimental rabbit exposed to ked bites. Green arrows point to *Anaplasma* sp., magnification x100.

This *Anaplasma* localized at the periphery of infected erythrocytes and appeared in dot-like forms similar to the morphology of *Anaplasma marginale*.

#### (i) Molecular analysis

DNA was isolated from the blood of the test animals and analyzed using PCR-HRM to detect the presence of *Anaplasma* infection (Fig 5).

**Fig 5:**
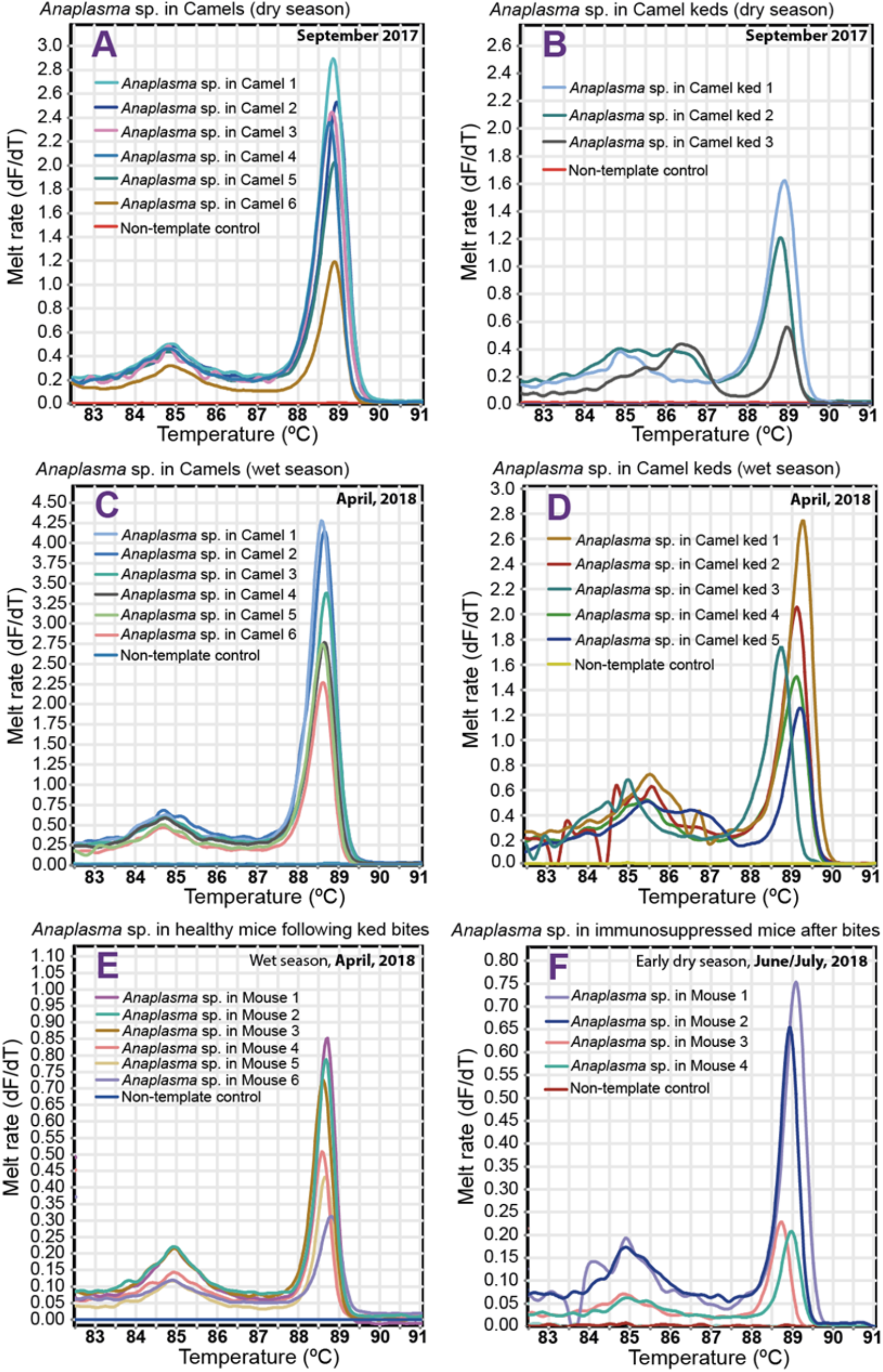
PCR - HRM melt curves for detection of *Anaplasma* sp. in mice, camels and keds. The experimental mice were exposed to the bites of freshly collected camel keds. Representative samples forming the peaks above were purified and sequenced to confirm identity (A - F). DNA sequencing has positively identified *Anaplasma* sp. in camels (A & C), keds (B & D), and mice (both healthy-E and immunosuppressed mice-F).

The prevalence of *Anaplasma* in the test mice for the three repeat experiments were 47.4%, 6.9% and 17.9%, respectively and 25% in rabbits (Table 2).

**Table 2:** Summary of *Anaplasma* infection in the test animals.

| Experiment | Season | Host | *Anaplasma* infection prevalence in test animals |
| --- | --- | --- | --- |
| 1<sup>st</sup> | Dry, September 2017 | Mice | 47.4% (n = 9/19) |
| 2<sup>nd</sup> | Wet, April 2018 | mice | 6.9% (n = 4/58) |
| 3<sup>rd</sup> | Late wet, July-August 2019 | Mice | 17.9% (n = 22/123) |
|  |  | Rabbits | 25% (n = 1/4) |

#### (ii) Identification of *Anaplasma* spp. in test animals by gene sequencing

PCR amplicons of representative positive test samples were sequenced using 1000-bp 16S rRNA and the shorter 300-bp 16S rRNA gene markers. Multiple sequence alignment of ∼1000-bp 16S rRNA *Anaplasmataceae* sequences obtained from camels and mouse shared 100% identity to ‘*Ca*. Anaplasma camelii’ (S1 Fig). Alignment of shorter *Anaplasmataceae*-specific 300-bp 16S rRNA sequences showed sequence identity of 100% between the sequences derived from experimental mice that were exposed to repeated ked bites. These sequences share 100% identity to ‘*Ca*. Anaplasma camelii’.

Phylogenetic investigation of sequences obtained from this study with related sequences from GenBank showed that they are genetically identical to ‘*Ca*. Anaplasma camelii’ previously sequenced from camels in Saudi Arabia and Iran and closely related to *Anaplasma platys* [10,38] (Fig 6).

**Fig 6:**
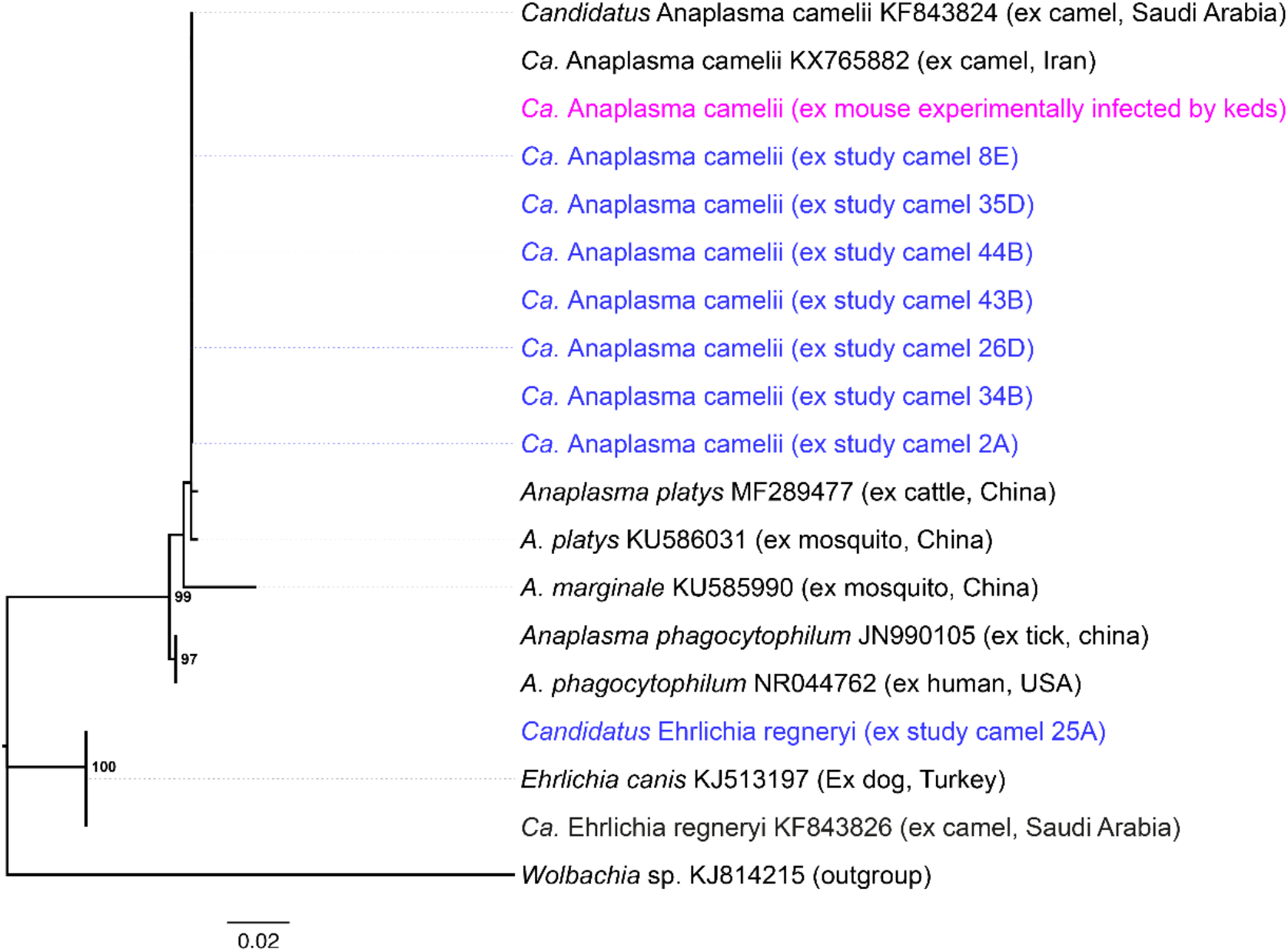
Phylogenetic tree of *Anaplasma* sp. based on full 16S rRNA nucleotide sequences. Tree was constructed with Neighbour-Joining method after aligning the sequences using MUSCLE in MEGA5. We used Kimura 2-parameter method with 1000 bootstrap repeats to calculate the distance matrices. The sequences obtained from this study are shown in bold font and number at the nodes represent the percentage bootstrap values. *Wolbachia* endosymbiont sequence (KJ814215) was used as outgroup.

### Ked-feeding bite exposure frequencies influenced transmission of *“Ca*. Anaplasma camelii”

The success of ‘*Ca*. Anaplasma camelii’ transmission to the experimental mice and rabbits by camel keds, *H. camelina*, during blood feeding was influenced by the exposure frequencies to fly bites (OR = 0.6504 (95% CI = 0.4714 – 0.8373), *p* = 0.0004, *n =* 115) (S1-S3 Tables). The pathogen transmission success could also be influenced by the proportion of keds that harbored *Anaplasma* (S5 Table). We observed that in some cases, a single exposure lasting for 12 h was sufficient to transmit *Anaplasma* to experimental mice (S3 and S4 Table), whilst in other instances, repeated exposure frequencies of up to 10 times did not transmit *Anaplasma* (S3 Table). Our repeat experiments to determine the vectorial competence of camel keds to transmit ‘*Ca*. Anaplasma camelii’ from infected camels to healthy mice (S1-S3 Tables), immunosuppressed mice (S2 Table), and rabbits (S3 Table) were successful.

## Discussion

This study reports the first experimental transmission of ‘*Ca*. Anaplasma camelii’ from naturally infected camels to laboratory mice and rabbits, demonstrating their vectorial competence in transmitting *Anaplasma*. Whilst ticks are required for cyclic transmission of *Anaplasma* [6], co-occurrent infestation with keds could imply direct involvement of these haematophagous flies in efficient transmission and maintenance of this haemopathogen within and between camel herds in northern Kenya. We recorded high ‘*Ca*. Anaplasma camelii’ infection rates of 63 – 78% in camels, and in contrast only 10 - 28.9% of camel keds carried *Ca*. Anaplasma camelii’ DNA during the various seasons. The presence of *Anaplasma* DNA in or on the camel keds was not surprising because when they bite infected camel to acquire bloodmeal, they also inadvertently ingest the accompanying blood-borne pathogens.

Field’s staining of thin blood smears in the present study revealed marginal occurrence of this pathogen in the host RBCs (Fig 4) similar to well-studied *Anaplasma marginale*. Further, this *Anaplasma* sp. was identified as ‘*Ca*. Anaplasma camelii’ by 16S rRNA gene sequencing. However, longer ∼1000-bp 16S rRNA gene sequences were impossible to get from keds, unlike in mice and camels, suggesting that ‘*Ca*. Anaplasma camelii’ does not multiply in the fly. This finding implies mechanical transmission of this *Anaplasma* species by the camel keds. Lower rates of *Anaplasma* prevalence in keds than in their camel hosts could suggest that in keds, ‘*Ca*. Anaplasma camelii’ does not transform or change in developmental forms, nor do they increase in numbers, thus over time they die off naturally and flies could acquire fresh bacteria through subsequent infected blood meals. These observations further support our hypothesis of mechanical transmission of *Anaplasma* sp. by keds, which often occurs via interrupted blood feeding. Further studies are needed to; (i) establish movement of ‘*Ca*. Anaplasma camelii’ in keds, and (ii) study the fate of this *Anaplasma* sp. in camels, keds, and in the test animals (mice and rabbits). The probability of *Anaplasma* transmission to the experimental animals by keds increased with the frequency fly bites. We also noted that the rates of contamination of keds with *Anaplasma* sp. varied in different geographical locations and thus the number of contaminated keds that successfully bite the host could also influence the chances of pathogen transmission. Transmission of *Anaplasma* was achieved in both healthy and immunosuppressed mice, as well as in rabbits. However, smaller percentage of immunosuppressed mice acquired *Anaplasma* infection possibly due to shortage of fly numbers in the wet season that permitted only a maximum of three ked-mice exposure frequencies.

16S rRNA *Anaplasma* sequences from camel, mouse, and rabbit showed 100% sequence identity as shown by multiple sequence alignment (S1 Fig). BLAST searches revealed 100% identity to ‘*Ca*. Anaplasma camelii’. This confirms common origin of “*Ca*. Anaplasma camelii” from camels and was subsequently transmitted to experimental mice and rabbits through ked feeding bites.

‘*Ca*. Anaplasma camelii’ species sequenced in this study was related to *Anaplasma platys*, a canine pathogen sequenced recently from cattle in China (GenBank accession number: MF289477). ‘*Ca*. Anaplasma camelii’ has not been characterized yet, thus poorly understood at present. Its host infection mechanism, target cells for infection, and pathogenicity in camels is presently not understood. Initial research studies using *in vitro* pathogen cultures in tick cell lines [39] will provide knowledge on the veterinary importance of this *Anaplasma* species.

The abundance of keds and their occurrence on vertebrate hosts throughout the year, in northern Kenya, enhances transmission and maintenance of *Anaplasma* pathogens in camels. Therefore, control of keds would be crucial in disease management and control for improved livestock productivity. We found the number of female camel keds to be almost twice that of males in random fly collections. Earlier studies on *Hippobosca equina* reported similar findings on sex ratios that were biased towards more females than males [40]. This is presumably important in maintaining ked populations to compensate for their low reproduction rates.

Further, our studies show that camel keds, *H. camelina*, are camel-specific because it largely infested camels, but hardly found on the other co-herded livestock species such as sheep, goats, cattle, donkeys, and dogs. Camels suffer the most burden from biting keds as they were infested by the highest numbers of flies than all the other co-herded livestock species, therefore making them most vulnerable to ked-borne diseases. Host selection by keds is possibly achieved through visual and olfactory signals as reported in other dipterans such as tsetse fly [41–44].

The highest ked count of up to 100 keds/camel occurred in the dry season in September 2017, whereas the lowest count of 7 keds/camel were recorded in the wet season in June-July 2018. Then, towards the late wet season, the fly numbers on camels gradually increased reaching highest numbers in September of the following dry season.

The number of camel infesting keds significantly reduced by 93% during the wet season and we observed that these flies leave their camel host to inhabit the surrounding vegetation in the grazing fields. Gradual rise in keds infesting camels was noted immediately at the end of rain season possibly indicating re-infestation of vertebrates by adult keds or their progeny. While seeking for preferred host for bloodmeal, these starved keds usually inflict bites on other presenting hosts, including humans.

Warm temperatures of about 32°C and relative humidity of 75% are key to successful pupal development in *Hippobosca equina* and *H. camelina* [45]. However, since wet seasons occur once or twice in a year in northern Kenya, it is reasonable that these keds, which are slow-breeders like their tsetse relatives, must continue to larviposit throughout the year to sustain their populations. Gravid females of *H. equina* deposit a single 3^rd^ instar larva every 3 - 8 days at larviposition sites established to be cracks in the mud walls of stables [45]. The present study aimed at determining the larviposition sites of gravid keds but without success. Better understanding of the life cycle and biology of camel keds is paramount for the design of fly control strategies. At present, there are no ked-specific strategies for controlling camel flies that are often assumed to be of little economic importance. For instance, there are no effective traps for these flies that infrequently depart from their camel host. Tsetse fly traps, such as NGU traps designed at *icipe* (Nairobi) work on the basis of visual (blue-black colour of the trap clothing) and olfactory cues (cow urine and acetone as attractants) can also trap biting flies such as *Stomoxys* and *Tabanus* species, but rarely camel keds as monoconical traps could only catch 0 – 3 keds per day/trap [21]. It is likely that keds perceive colours differently and olfactory cues, unlike tsetse fly, their closest relatives – hippoboscids and tsetse flies belong to the same superfamily, *Hippoboscoidae*. It is important to understand how camel-specific *H. camelina* recognizes its camel host via visual and olfactory cues in order to model ked-specific traps.

All camel keepers whose camels were sampled in the present study also kept domestic animals including dogs and small ruminants. Co-herding of livestock and domestic animals promotes spread of vector-borne diseases. For instance, *Anaplasma platys* infection in dogs could be transmitted to camels [19] or to humans [13–15]. *A. phagocytophilum* that commonly affects cattle is also known to be pathogenic to humans causing Human Granulocytic Anaplasmosis [46–48]. ‘*Ca*. Anaplasma camelii’ was recently reported in dromedary camels in Saudi Arabia [10], Iran [38], and Morocco [12]. Tick vectors belonging to genus *Hyalomma* were suspected to transmit ‘*Ca*. Anaplasma camelii’ in Moroccan one-humped camels that are reportedly the reservoir host of *Anaplasma* spp. [12].

Our study was limited by short sampling periods lasting between 7 – 30 days that limited pathogen transmission studies that relied on the field-collected keds, as we could not establish long-term laboratory colonies of flies. This limited frequencies of ked bites on the test animals, possibly reducing pathogen infection rates. Additionally, seasonal variation of ked populations also affected the transmission experiment. For instance, during the wet season in April, the ked numbers were very low and this limited our experiments.

In conclusion, we also demonstrate for the first time the ability of camel-specific keds, *Hippobosca camelina*, to transmit ‘*Ca*. Anaplasma camelii’ originating from naturally infected dromedary camels into laboratory-reared mice and rabbits through contaminated ked bites. The prevalence of camel anaplasmosis caused by a single *Anaplasma* sp. was high throughout the seasons, unlike in keds with low rates of contamination but still achieved to efficiently transmit pathogens in all our test groups of animals. Further research studies are needed to determine the vectorial competence of keds in transmission of other blood-borne pathogens of veterinary and zoonotic importance.

## Acknowledgements

We are grateful to all camel farmers from the study area for allowing us to collect samples. Many thanks to Galtumo Lordagos, Ogoga Kaldale, Lmachungwan Galgitele, Ltulusuan Letoiye, Moika Naparakwo, Fereiti Bargul, and Ali Baltor for assisting in restraining camels during collection of blood and ked samples. John Ng’iela of *icipe*’s Animal Health Department is acknowledged for providing valuable support during field sampling, staining and interpreting Field’s stained blood smears for identification of *Anaplasma* sp. Winny Cherono, currently a graduate member at the Institute of Surveyors of Kenya (GIS Chapter), generated maps for this study. We are also very grateful to Collins Kigen for his technical support.

## Author contributions

Conceived and designed the experiments: JLB, KK, RSC, MC, DKM Performed the experiments: JLB, KK, RSC

Analyzed the data: JLB, KK, MG, JV, RSC, JMM, MC, DKM Contributed reagents/materials/analysis tools: JLB, MG, JV, MC, DKM Wrote the first draft: JLB

All authors reviewed, edited and approved the final manuscript.

