## Supplementary Table 1 for "Transmission of ‘*Candidatus* Anaplasma camelii’ to laboratory animals by camel-specific keds, *Hippobosca camelina*"

**S1 Table: Showing feeding schedule of camel keds on healthy Swiss white mice for pathogen transmission experiment.** Detection of *“Ca.* Anaplasma camelii*”* in experimental mice group was determined by PCR-HRM using genus-specific primers for 16S rRNA gene target. The data shows that 47.4% of mice in the test group (*n* = 9/19) have acquired *Anaplasma* infection following ked feeding bites. The control mice group (n = 2) were not exposed to biting flies.

| **S/No.** | **Mouse ID** | **Total keds**  (20 flies/mouse) | **Dates exposed**  (April 2018) | **Exposure frequency** | ***Anaplasma* infection status** |
| --- | --- | --- | --- | --- | --- |
| 1 | 1A (control 1) | Not applicable (N/A) | N/A | N/A | Negative |
| 2 | 1B (control 2) | N/A | N/A | N/A | Negative |
| 3 | 2A | 100 | 7^th^, 8^th^, 11^th^, 14^th^, 21^st^ | 5 | Negative |
| 4 | 2B | 100 | 7^th^, 8^th^, 11^th^, 14^th^, 21^st^ | 5 | *Anaplasma* sp. |
| 5 | 3A | 120 | 5^th^, 6^th^, 9^th^, 11^th^, 17^th^, 20^th^ | 6 | *Anaplasma* sp. |
| 6 | 3B | 120 | 5^th^, 6^th^, 9^th^, 11^th^, 17^th^, 20^th^ | 6 | Negative |
| 7 | 3C | 120 | 5^th^, 6^th^, 9^th^, 11^th^, 17^th^, 20^th^ | 6 | *Anaplasma* sp. |
| 8 | 4A | 100 | 10^th^, 12^th^, 17^th^, 18^th^, 21^st^ | 5 | Negative |
| 9 | 4B | 100 | 10^th^, 12^th^, 17^th^, 18^th^, 21^st^ | 5 | Negative |
| 10 | 5A | 100 | 5^th^, 6^th^, 9^th^, 12^th^, 15th | 5 | Negative |
| 11 | 5B | 100 | 5^th^, 6^th^, 9^th^, 12^th^, 15^th^ | 5 | Negative |
| 12 | 6A | 100 | 7^th^, 8^th^, 10^th^, 12^th^, 16^th^ | 5 | Negative |
| 13 | 6B | 100 | 7^th^, 8^th^, 10^th^, 12^th^, 16^th^ | 5 | Negative |
| 14 | 6C | 100 | 7^th^, 8^th^, 10^th^, 12^th^, 16^th^ | 5 | *Anaplasma* sp. |
| 15 | 7A | 60 | 11^th^, 13^th^, 16 | 3 | Negative |
| 16 | 7B | 60 | 11^th^, 13^th^, 16 | 3 | *Anaplasma* sp. |
| 17 | 8 | 40 | 11^th^, 15^th^ | 2 | *Anaplasma* sp. |
| 18 | 9A | 20 | 18^th^ | 1 | *Anaplasma* sp. |
| 19 | 9B | 20 | 18^th^ | 1 | Negative |
| 20 | 10A | 20 | 19^th^ | 1 | *Anaplasma* sp. |
| 21 | 10B | 20 | 19^th^ | 1 | *Anaplasma* sp. |
