## Supplementary Table 2 for "Transmission of ‘*Candidatus* Anaplasma camelii’ to laboratory animals by camel-specific keds, *Hippobosca camelina*"

**S2 Table: Feeding schedule of camel keds on caesium-immunosuppressed Swiss white mice (*n* = 60) for *Anaplasma* transmission to determine the role of immunity during infection.** Overall, we detected *Anaplasma* infection in 6.9% of test mice (*n* = 4/58). PCR-HRM. Control mice were not exposed to biting flies. We recorded *Anaplasma* infection of 12.9% (*n* = 4/31) in mice screened after 60 days, but all 27 mice that were sacrificed 140 days post-ked exposure did not have infection.

| S/No. | Mouse ID | Total keds  (20 flies/mouse) | Date(s) exposed | Exposure frequency | *Anaplasma* infection status |
| --- | --- | --- | --- | --- | --- |
| 1 | Control 1 | Not applicable (N/A) | N/A | N/A | Negative |
| 2 | Control 2 | N/A | N/A | N/A | Negative |
| 3 | 1A | 60 | 28.06.18, 3.07.18, 4.07.18 | 3 | Negative |
| 4 | 1B | 60 | 28.06.18, 3.07.18, 4.07.18 | 3 | Negative |
| 5 | 1C | 60 | 28.06.18, 3.07.18, 4.07.18 | 3 | Negative |
| 6 | 1D | 60 | 28.06.18, 3.07.18, 4.07.18 | 3 | Negative |
| 7 | 1E | 60 | 28.06.18, 3.07.18, 4.07.18 | 3 | Negative |
| 8 | 1F | 60 | 28.06.18, 3.07.18, 4.07.18 | 3 | *Anaplasma* sp. |
| 9 | 1G | 60 | 28.06.18, 3.07.18, 4.07.18 | 3 | Negative |
| 10 | 1H | 60 | 28.06.18, 3.07.18, 4.07.18 | 3 | Negative |
| 11 | 1I | 60 | 28.06.18, 3.07.18, 4.07.18 | 3 | Negative |
| 12 | 1J | 60 | 28.06.18, 3.07.18, 4.07.18 | 3 | Negative |
| 13 | 1K | 60 | 28.06.18, 3.07.18, 4.07.18 | 3 | Negative |
| 14 | 1L | 60 | 28.06.18, 3.07.18, 4.07.18 | 3 | Negative |
| 14 | 2A | 20 | 29.06.2018 | 1 | Negative |
| 16 | 2B | 20 | 29.06.2018 | 1 | Negative |
| 17 | 2C | 20 | 29.06.2018 | 1 | *Anaplasma* sp. |
| 18 | 2D | 20 | 29.06.2018 | 1 | Negative |
| 19 | 2E | 20 | 29.06.2018 | 1 | Negative |
| 20 | 3A | 40 | 27.06.18 & 4.07.18 | 2 | Negative |
| 21 | 3B | 40 | 27.06.18 & 4.07.18 | 2 | Negative |
| 22 | 3C | 40 | 27.06.18 & 4.07.18 | 2 | Negative |
| 23 | 3D | 40 | 27.06.18 & 4.07.18 | 2 | *Anaplasma* sp. |
| 24 | 4A | 20 | 29.06.2018 | 1 | Negative |
| 25 | 4B | 20 | 29.06.2018 | 1 | *Anaplasma* sp. |
| 26 | 4C | 20 | 29.06.2018 | 1 | Negative |
| 27 | 4D | 20 | 29.06.2018 | 1 | Negative |
| 28 | 4E | 20 | 29.06.2018 | 1 | Negative |
| 29 | 4F | 20 | 29.06.2018 | 1 | Negative |
| 30 | 4G | 20 | 29.06.2018 | 1 | Negative |
| 31 | 4H | 20 | 29.06.2018 | 1 | Negative |
| 32 | 4I | 20 | 29.06.2018 | 1 | Negative |
| 33 | 5A | 20 | 26.06.2018 | 1 | Negative |
| 34 | 5B | 20 | 26.06.2018 | 1 | Negative |
| 35 | 6A | 20 | 26.06.2018 | 1 | Negative |
| 36 | 6B | 20 | 26.06.2018 | 1 | Negative |
| 37 | 6C | 20 | 26.06.2018 | 1 | Negative |
| 38 | 6D | 20 | 26.06.2018 | 1 | Negative |
| 39 | 6E | 20 | 26.06.2018 | 1 | Negative |
| 40 | 6F | 20 | 26.06.2018 | 1 | Negative |
| 41 | 6G | 20 | 26.06.2018 | 1 | Negative |
| 42 | 6H | 20 | 26.06.2018 | 1 | Negative |
| 43 | 6I | 20 | 26.06.2018 | 1 | Negative |
| 44 | 7A | 20 | 3.07.2018 | 1 | Negative |
| 45 | 7B | 20 | 3.07.2018 | 1 | Negative |
| 46 | 7C | 20 | 3.07.2018 | 1 | Negative |
| 47 | 7D | 20 | 3.07.2018 | 1 | Negative |
| 48 | 7E | 20 | 3.07.2018 | 1 | Negative |
| 49 | 7F | 20 | 3.07.2018 | 1 | Negative |
| 50 | 7G | 20 | 3.07.2018 | 1 | Negative |
| 51 | 7H | 20 | 3.07.18 | 1 | Negative |
| 52 | 8A | 60 | 4.07.18, 8.07.18, 9.07.18 | 3 | Negative |
| 53 | 8B | 60 | 4.07.18, 8.07.18, 9.07.18 | 3 | Negative |
| 54 | 8C | 60 | 4.07.18, 8.07.18, 9.07.18 | 3 | Negative |
| 55 | 8D | 60 | 4.07.18, 8.07.18, 9.07.18 | 3 | Negative |
| 56 | 9A | 20 | 4.07.18, 8.07.18, 9.07.18 | 1 | Negative |
| 57 | 9B | 20 | 4.07.18, 8.07.18, 9.07.18 | 1 | Negative |
| 58 | 9C | 20 | 4.07.18, 8.07.18, 9.07.18 | 1 | Negative |
| 59 | 9D | 20 | 4.07.18, 8.07.18, 9.07.18 | 1 | Negative |
| 60 | 9E |  | 4.07.18, 8.07.18, 9.07.18 | 1 | Negative |
