## Supplementary Table 3 for "Transmission of ‘*Candidatus* Anaplasma camelii’ to laboratory animals by camel-specific keds, *Hippobosca camelina*"

**S3 Table: Feeding schedule of ex-camel keds on mice (n = 123) and rabbits (n = 6) for *Anaplasma* transmission study to determine vectorial competence.** PCR-HRM analysis using genus-specific 16S rRNA gene target detected *Anaplasma* infection in 17.9% of test mice (n = 22/123) and in 25% of rabbits (n = 1/4) post-ked bites. Control mice (n = 8) and rabbits (n = 2) were not exposed to any fly bites.

| **S/No.** | **Mouse ID** | **Total exposed keds**  (20 flies/mouse) | **Dates exposed**  (July-August 2019) | **Exposure frequency** | ***Anaplasma* infection status** |
| --- | --- | --- | --- | --- | --- |
| 1. **Mice experiments:** | | | | | |
| 1 | 1A | Control | Not exposed to biting flies including keds | 0 | Negative |
| 2 | 1B | Control |  | 0 | Negative |
| 3 | 1C | Control |  | 0 | Negative |
| 4 | 1D | Control |  | 0 | Negative |
| 5 | 1E | Control |  | 0 | Negative |
| 6 | 1F | Control |  | 0 | Negative |
| 7 | 1G | Control |  | 0 | Negative |
| 8 | 1F | Control |  | 0 | Negative |
| 9 | 2A | 80 | 7^th^ July, 15^th^ July, 16^th^ July, 18^th^ July | 4 | Negative |
| 10 | 2B | 80 |  | 4 | *Anaplasma* sp*.* |
| 11 | 2C | 80 |  | 4 | *Anaplasma* sp*.* |
| 12 | 2D | 80 |  | 4 | *Anaplasma* sp*.* |
| 13 | 2E | 80 |  | 4 | *Anaplasma* sp*.* |
| 14 | 2F | 80 |  | 4 | *Anaplasma* sp. |
| 15 | 2G | 80 |  | 4 | *Anaplasma* sp. |
| 16 | 2H | 80 |  | 4 | Negative |
| 17 | 2I | 80 |  | 4 | *Anaplasma* sp. |
| 18 | 2J | 80 |  | 4 | Negative |
| 19 | 3A | 140 | 8^th^ July, 10^th^ July, 15^th^ July, 16^th^ July, 24^th^ July, 2^nd^ Aug, 3^rd^ Aug | 7 | *Anaplasma* sp. |
| 20 | 3B | 140 |  | 7 | Negative |
| 21 | 3C | 140 |  | 7 | Negative |
| 22 | 3D | 140 |  | 7 | Negative |
| 23 | 3E | 140 |  | 7 | Negative |
| 24 | 3F | 140 |  | 7 | Negative |
| 25 | 3G | 140 |  | 7 | Negative |
| 26 | 3H | 140 |  | 7 | Negative |
| 27 | 3I | 140 |  | 7 | Negative |
| 28 | 3J | 140 |  | 7 | Negative |
| 29 | 3K | 140 |  | 7 | Negative |
| 30 | 3L | 140 |  | 7 | Negative |
| 31 | 3M | 140 |  | 7 | Negative |
| 32 | 3N | 140 |  | 7 | Negative |
| 33 | 3O | 140 |  | 7 | Negative |
| 34 | 3P | 140 |  | 7 | Negative |
| 35 | 4A | 120 | 6^th^ July, 11^th^ July, 13^th^ July, 14^th^ July, 17^th^ July, 23^rd^ July | 6 | *Anaplasma* sp. |
| 36 | 4B | 120 |  | 6 | Negative |
| 37 | 4C | 120 |  | 6 | *Anaplasma* sp. |
| 38 | 4D | 120 |  | 6 | *Anaplasma* sp. |
| 39 | 4E | 120 |  | 6 | *Anaplasma* sp. |
| 40 | 4F | 120 |  | 6 | Negative |
| 41 | 4G | 120 |  | 6 | Negative |
| 42 | 4H | 120 |  | 6 | *Anaplasma* sp. |
| 43 | 4I | 120 |  | 6 | Negative |
| 44 | 4J | 120 |  | 6 | *Anaplasma* sp. |
| 45 | 5A | 80 | 23^rd^ July, 29^th^ July, 30^th^ July, 2^nd^ Aug | 4 | *Anaplasma* sp. |
| 46 | 5B | 80 |  | 4 | *Anaplasma* sp. |
| 47 | 5C | 80 |  | 4 | Negative |
| 48 | 5D | 80 |  | 4 | Negative |
| 49 | 5E | 80 |  | 4 | Negative |
| 50 | 5F | 80 |  | 4 | Negative |
| 51 | 5G | 80 |  | 4 | Negative |
| 52 | 5H | 80 |  | 4 | Negative |
| 53 | 6A | 80 | 8^th^ July, 28^th^ July, 29^th^ July, 30^th^ July | 4 | Negative |
| 54 | 6B | 80 |  | 4 | Negative |
| 55 | 6C | 80 |  | 4 | Negative |
| 56 | 6D | 80 |  | 4 | *Anaplasma* sp. |
| 57 | 6E | 80 |  | 4 | Negative |
| 58 | 6F | 80 |  | 4 | Negative |
| 59 | 6G | 80 |  | 4 | Negative |
| 60 | 6H | 80 |  | 4 | Negative |
| 61 | 6I | 80 |  | 4 | *Anaplasma* sp. |
| 62 | 7A | 80 | 23^rd^ July, 29^th^ July, 30^th^ July, 2^nd^ Aug | 4 | Negative |
| 63 | 7B | 80 |  | 4 | Negative |
| 64 | 7C | 80 |  | 4 | Negative |
| 65 | 7D | 80 |  | 4 | Negative |
| 66 | 7E | 80 |  | 4 | Negative |
| 67 | 7F | 80 |  | 4 | Negative |
| 68 | 7G | 80 |  | 4 | Negative |
| 69 | 7H | 80 |  | 4 | Negative |
| 70 | 8A | 180 | 15^th^ July, 17^th^ July, 18^th^ July, 19^th^ July, 20^th^ July, 21^st^ July, 28^th^ July, 29^th^ July, 30^th^ July | 9 | Negative |
| 71 | 8B | 180 |  | 9 | Negative |
| 72 | 8C | 180 |  | 9 | Negative |
| 73 | 8D | 180 |  | 9 | Negative |
| 74 | 8E | 180 |  | 9 | Negative |
| 75 | 8F | 180 |  | 9 | Negative |
| 76 | 8G | 180 |  | 9 | Negative |
| 77 | 8H | 180 |  | 9 | Negative |
| 78 | 8I | 180 |  | 9 | Negative |
| 79 | 8J | 180 |  | 9 | Negative |
| 80 | 8K | 180 |  | 9 | Negative |
| 81 | 8L | 180 |  | 9 | Negative |
| 82 | 8M | 180 |  | 9 | Negative |
| 83 | 8N | 180 |  | 9 | Negative |
| 84 | 8O | 180 |  | 9 | Negative |
| 85 | 8P | 180 |  | 9 | Negative |
| 86 | 9A | 200 | 7^th^ July, 8^th^ July, 15^th^ July, 18^th^ July, 19^th^ July, 20^th^ July, 21^st^ July, 25^th^ July, 29^th^ July, 30^th^ July | 10 | Negative |
| 87 | 9B | 200 |  | 10 | Negative |
| 88 | 9C | 200 |  | 10 | Negative |
| 89 | 9D | 200 |  | 10 | Negative |
| 90 | 9E | 200 |  | 10 | Negative |
| 91 | 9F | 200 |  | 10 | Negative |
| 92 | 9G | 200 |  | 10 | Negative |
| 93 | 9H | 200 |  | 10 | Negative |
| 94 | 9I | 200 |  | 10 | Negative |
| 95 | 9J | 200 |  | 10 | Negative |
| 96 | 9K | 200 |  | 10 | Negative |
| 97 | 9L | 200 |  | 10 | Negative |
| 98 | 9M | 200 |  | 10 | Negative |
| 99 | 9N | 200 |  | 10 | Negative |
| 100 | 9O | 200 |  | 10 | Negative |
| 101 | 9P | 200 |  | 10 | Negative |
| 102 | 9Q | 200 |  | 10 | Negative |
| 103 | 9R | 200 |  | 10 | Negative |
| 104 | 10A | 80 | 25^th^ July, 28^th^ July, 29^th^ July, 30^th^ July | 4 | *Anaplasma* sp. |
| 105 | 10B | 80 |  | 4 | Negative |
| 106 | 10C | 80 |  | 4 | *Anaplasma* sp. |
| 107 | 10D | 80 |  | 4 | Negative |
| 108 | 10E | 80 |  | 4 | Negative |
| 109 | 10F | 80 |  | 4 | Negative |
| 110 | 10G | 80 |  | 4 | Negative |
| 111 | 10H | 80 |  | 4 | Negative |
| 112 | 10I | 80 |  | 4 | Negative |
| 113 | 10J | 80 |  | 4 | Negative |
| 114 | 10K | 80 |  | 4 | Negative |
| 115 | 10L | 80 |  | 4 | Negative |
| 116 | 10M | 80 |  | 4 | Negative |
| 117 | 10N | 80 |  | 4 | Negative |
| 118 | 100 | 80 |  | 4 | Negative |
| 119 | 10P | 80 |  | 4 | Negative |
| 120 | 10Q | 80 |  | 4 | Negative |
| 121 | 10R | 80 |  | 4 | *Anaplasma* sp. |
| 122 | 10S | 80 |  | 4 | Negative |
| 123 | 10T | 80 |  | 4 | *Anaplasma* sp. |
| 2. **Rabbit experiments:** | | | | | |
| 1 | R1 | 120 | 10^th^ July, 13^th^ July, 16^th^ July, 20^th^ July, 22^nd^ July, 26^th^ July | 6 | Negative |
| 2 | R2 | 140 | 13^th^ July, 14^th^ July, 15^th^ July, 16^th^ July, 20^th^ July, 22^nd^ July, 26^th^ July | 7 | *Anaplasma* sp*.* |
| 3 | R3 | 140 | 13th July, 14th July, 15th July, 16th July, 20th July, 22nd July, 26th July | 7 | Negative |
| 4 | R4 | 140 | 13th July, 14th July, 15th July, 16th July, 20th July, 22nd July, 26th July | 7 | Negative |
| 5 | Jane | Control | Not exposed to biting flies including keds | 0 | Negative |
| 6 | Jasmin | Control |  | 0 | Negative |

The mice were divided into 10 groups; group one are the control mice that were not exposed to ked bites, whereas group 2 – 10 were the test mice which were exposed to ked bites at varying frequencies on different dates. R1 – R4 are the test rabbits, whereas rabbits named Jane and Jasmin belonged to the control group.
