## Supplementary Table 4 for "Transmission of ‘*Candidatus* Anaplasma camelii’ to laboratory animals by camel-specific keds, *Hippobosca camelina*"

**S4 Table: Seasonal variation in prevalence of “*Ca.* Anaplasma camelii” in camels and camel keds, *H. camelina*.**

| Sampling site | Date | Season | Host | Prevalence of *Anaplasma* sp. |
| --- | --- | --- | --- | --- |
| Koya  (N 01° 23' 11";  E 37° 57' 11.7") | September 2017 | Dry | Camels | 70.3% (175/249) |
|  |  |  | *H. camelina* | 9.9% (19/192) |
| Laisamis town  (N 01° 35' 16.1";  E 037° 48' 26.2") | June-July 2018 | Wet | Camels | 63.9% (179/280) |
|  |  |  | *H. camelina* | 20.8% (20/96) |
| Silapani  (N 01° 39’ 25.7”;  E 037° 49’ 00.6”) | July-August 2019 | Late wet | Camels | 77.9% (348/447) |
|  |  |  | *H. camelina* | 28.9% (22/76) |
