## Supplementary Table 5 for "Transmission of ‘*Candidatus* Anaplasma camelii’ to laboratory animals by camel-specific keds, *Hippobosca camelina*"

**S5 Table: Contamination rates of camel keds collected from camel herds occupying various geographical locations in Laisamis sub-County.**

| **Site** | **GPS coordinate** | **Year of sampling** | **Prevalence of *Anaplasma* in keds** |
| --- | --- | --- | --- |
| Silapani | N01° 39’ 25.7”  E037° 49’ 00.6” | 2019 | 14.85% (15/101) |
| Silapani | N01° 39’ 25.7”  E037° 49’ 00.6” | 2019 | 16.13% (5/31) |
| Silapani | N01° 39’ 25.7”  E037° 49’ 00.6” | 2019 | 16% (12/75) |
| Silapani | N01° 39’ 25.7”  E037° 49’ 00.6” | 2019 | 21.74% (5/23) |
| Sakardala -polytechnique | N01° 37’ 26.2”  E037° 48’ 31.6’’ | 2019 | 18.75% (6/32) |
| Nkwanja e Ndege |  | 2019 | 17.86% (10/56) |
| Kargi | N02° 31’ 0”  E037° 34’ 0” | 2019 | 8.7% (6/69) |
| Sirata | N01° 38’ 02.5”  E037° 48’ 22.7” | 2019 | 15% (3/20)  9.68% (3/31) |
| Sirata | N01° 38’ 02.5”  E037° 48’ 22.7 | 2018 | 33.33% (4/12) |
| Nkwanja e Ndege |  | 2018 | 19.35% (6/31) |
| Manyatta Secondary | N01° 34’ 48.9”  E037° 47’ 38.8’’ | 2018 | 32.14% (9/28) |
| Soweto | N01° 35’ 42.8”  E037° 48’ 40.4’’ | 2018 | 34.38% (11/32) |
| Tirgamo | N01° 35’ 39.8”  E037° 54’ 11.6’’ | 2018 | 25% (3/12) |
| Nkang e Lawai |  | 2018 | 25% (3/12) |
