## Supplementary Figure 1 for "Transmission of ‘*Candidatus* Anaplasma camelii’ to laboratory animals by camel-specific keds, *Hippobosca camelina*"

```
3A_M__C         -------ATTCGTGCCAGCAGCCGCGGTAATACGGAGGGGGCAAGCGTTGTTCGGAATTA
8_E             -----------------------------ATACGGAGGGGGCAAGCGTTGTTCGGAATTA
2_A             -----------------------------------AGGGGGCAAGCGTTGTTCGGAATTA
43_B            --------TCCGTGCCAGCAGCCGCGGTAATACGGAGGGGGCAAGCGTTGTTCGGAATTA
34_B            CGGCAAACTCCGTGCCAGCAGCCGCGGTAATACGGAGGGGGCAAGCGTTGTTCGGAATTA
45_B            CGGCAAACTCCGTGCCAGCAGCCGCGGTAATACGGAGGGGGCAAGCGTTGTTCGGAATTA
26_D            CGGCAAACTCCGTGCCAGCAGCCGCGGTAATACGGAGGGGGCAAGCGTTGTTCGGAATTA
35_D            CGGCAAACTCCGTGCCAGCAGCCGCGGTAATACGGAGGGGGCAAGCGTTGTTCGGAATTA
                                                   *************************

3A_M__C         TTGGGCGTAAAGGGCATGTAGGCGGTTCGGTAAGTTAAAGGTGAAATGCCAGGGCTTAAC
8_E             TTGGGCGTAAAGGGCATGTAGGCGGTTCGGTAAGTTAAAGGTGAAATGCCAGGGCTTAAC
2_A             TTGGGCGTAAAGGGCATGTAGGCGGTTCGGTAAGTTAAAGGTGAAATGCCAGGGCTTAAC
43_B            TTGGGCGTAAAGGGCATGTAGGCGGTTCGGTAAGTTAAAGGTGAAATGCCAGGGCTTAAC
34_B            TTGGGCGTAAAGGGCATGTAGGCGGTTCGGTAAGTTAAAGGTGAAATGCCAGGGCTTAAC
45_B            TTGGGCGTAAAGGGCATGTAGGCGGTTCGGTAAGTTAAAGGTGAAATGCCAGGGCTTAAC
26_D            TTGGGCGTAAAGGGCATGTAGGCGGTTCGGTAAGTTAAAGGTGAAATGCCAGGGCTTAAC
35_D            TTGGGCGTAAAGGGCATGTAGGCGGTTCGGTAAGTTAAAGGTGAAATGCCAGGGCTTAAC
                ************************************************************

3A_M__C         CCTGGAGCTGCTTTTAATACTGCCAGACTCGAGTCCGGGAGAGGATAGCGGAATTCCTAG
8_E             CCTGGAGCTGCTTTTAATACTGCCAGACTCGAGTCCGGGAGAGGATAGCGGAATTCCTAG
2_A             CCTGGAGCTGCTTTTAATACTGCCAGACTCGAGTCCGGGAGAGGATAGCGGAATTCCTAG
43_B            CCTGGAGCTGCTTTTAATACTGCCAGACTCGAGTCCGGGAGAGGATAGCGGAATTCCTAG
34_B            CCTGGAGCTGCTTTTAATACTGCCAGACTCGAGTCCGGGAGAGGATAGCGGAATTCCTAG
45_B            CCTGGAGCTGCTTTTAATACTGCCAGACTCGAGTCCGGGAGAGGATAGCGGAATTCCTAG
26_D            CCTGGAGCTGCTTTTAATACTGCCAGACTCGAGTCCGGGAGAGGATAGCGGAATTCCTAG
35_D            CCTGGAGCTGCTTTTAATACTGCCAGACTCGAGTCCGGGAGAGGATAGCGGAATTCCTAG
                ************************************************************

3A_M__C         TGTAGAGGTGAAATTCGTAGATATTAGGAGGAACACCAGTGGCGAAGGCGGCTATCTGGT
8_E             TGTAGAGGTGAAATTCGTAGATATTAGGAGGAACACCAGTGGCGAAGGCGGCTATCTGGT
2_A             TGTAGAGGTGAAATTCGTAGATATTAGGAGGAACACCAGTGGCGAAGGCGGCTATCTGGT
43_B            TGTAGAGGTGAAATTCGTAGATATTAGGAGGAACACCAGTGGCGAAGGCGGCTATCTGGT
34_B            TGTAGAGGTGAAATTCGTAGATATTAGGAGGAACACCAGTGGCGAAGGCGGCTATCTGGT
45_B            TGTAGAGGTGAAATTCGTAGATATTAGGAGGAACACCAGTGGCGAAGGCGGCTATCTGGT
26_D            TGTAGAGGTGAAATTCGTAGATATTAGGAGGAACACCAGTGGCGAAGGCGGCTATCTGGT
35_D            TGTAGAGGTGAAATTCGTAGATATTAGGAGGAACACCAGTGGCGAAGGCGGCTATCTGGT
                ************************************************************

3A_M__C         CCGGTACTGACGCTGAGGTGCGAAAGCGTGGGGAGCAAACAGGATTAGATACCCTGGTAG
8_E             CCGGTACTGACGCTGAGGTGCGAAAGCGTGGGGAGCAAACAGGATTAGATACCCTGGTAG
2_A             CCGGTACTGACGCTGAGGTGCGAAAGCGTGGGGAGCAAACAGGATTAGATACCCTGGTAG
43_B            CCGGTACTGACGCTGAGGTGCGAAAGCGTGGGGAGCAAACAGGATTAGATACCCTGGTAG
34_B            CCGGTACTGACGCTGAGGTGCGAAAGCGTGGGGAGCAAACAGGATTAGATACCCTGGTAG
45_B            CCGGTACTGACGCTGAGGTGCGAAAGCGTGGGGAGCAAACAGGATTAGATACCCTGGTAG
26_D            CCGGTACTGACGCTGAGGTGCGAAAGCGTGGGGAGCAAACAGGATTAGATACCCTGGTAG
35_D            CCGGTACTGACGCTGAGGTGCGAAAGCGTGGGGAGCAAACAGGATTAGATACCCTGGTAG
                ************************************************************

3A_M__C         TCCACGCTGTAAACGATGAGTGCTGAATGTGGGGACGTTTTGTCTCTGTGTTGTAGCTAA
8_E             TCCACGCTGTAAACGATGAGTGCTGAATGTGGGGACGTTTTGTCTCTGTGTTGTAGCTAA
2_A             TCCACGCTGTAAACGATGAGTGCTGAATGTGGGGACGTTTTGTCTCTGTGTTGTAGCTAA
43_B            TCCACGCTGTAAACGATGAGTGCTGAATGTGGGGACGTTTTGTCTCTGTGTTGTAGCTAA
34_B            TCCACGCTGTAAACGATGAGTGCTGAATGTGGGGACGTTTTGTCTCTGTGTTGTAGCTAA
45_B            TCCACGCTGTAAACGATGAGTGCTGAATGTGGGGACGTTTTGTCTCTGTGTTGTAGCTAA
26_D            TCCACGCTGTAAACGATGAGTGCTGAATGTGGGGACGTTTTGTCTCTGTGTTGTAGCTAA
35_D            TCCACGCTGTAAACGATGAGTGCTGAATGTGGGGACGTTTTGTCTCTGTGTTGTAGCTAA
                ************************************************************

3A_M__C         CGCGTTAAGCACTCCGCCTGGGGACTACGGTCGCAAGACTAAAACTCAAAGGAATTGACG
8_E             CGCGTTAAGCACTCCGCCTGGGGACTACGGTCGCAAGACTAAAACTCAAAGGAATTGACG
2_A             CGCGTTAAGCACTCCGCCTGGGGACTACGGTCGCAAGACTAAAACTCAAAGGAATTGACG
43_B            CGCGTTAAGCACTCCGCCTGGGGACTACGGTCGCAAGACTAAAACTCAAAGGAATTGACG
34_B            CGCGTTAAGCACTCCGCCTGGGGACTACGGTCGCAAGACTAAAACTCAAAGGAATTGACG
45_B            CGCGTTAAGCACTCCGCCTGGGGACTACGGTCGCAAGACTAAAACTCAAAGGAATTGACG
26_D            CGCGTTAAGCACTCCGCCTGGGGACTACGGTCGCAAGACTAAAACTCAAAGGAATTGACG
35_D            CGCGTTAAGCACTCCGCCTGGGGACTACGGTCGCAAGACTAAAACTCAAAGGAATTGACG
                ************************************************************

3A_M__C         GGGACCCGCACAAGCGGTGGAGCATGTGGTTTAATTCGATGCAACGCGAAGAACCTTACC
8_E             GGGACCCGCACAAGCGGTGGAGCATGTGGTTTAATTCGATGCAACGCGAAGAACCTTACC
2_A             GGGACCCGCACAAGCGGTGGAGCATGTGGTTTAATTCGATGCAACGCGAAGAACCTTACC
43_B            GGGACCCGCACAAGCGGTGGAGCATGTGGTTTAATTCGATGCAACGCGAAGAACCTTACC
34_B            GGGACCCGCACAAGCGGTGGAGCATGTGGTTTAATTCGATGCAACGCGAAGAACCTTACC
45_B            GGGACCCGCACAAGCGGTGGAGCATGTGGTTTAATTCGATGCAACGCGAAGAACCTTACC
26_D            GGGACCCGCACAAGCGGTGGAGCATGTGGTTTAATTCGATGCAACGCGAAGAACCTTACC
35_D            GGGACCCGCACAAGCGGTGGAGCATGTGGTTTAATTCGATGCAACGCGAAGAACCTTACC
                ************************************************************

3A_M__C         ACTTCTTGACATGGAGATTAGATCCTTCTTAACAGAAGGGCGCAGTTCGGCTGGATCTCG
8_E             ACTTCTTGACATGGAGATTAGATCCTTCTTAACAGAAGGGCGCAGTTCGGCTGGATCTCG
2_A             ACTTCTTGACATGGAGATTAGATCCTTCTTAACAGAAGGGCGCAGTTCGGCTGGATCTCG
43_B            ACTTCTTGACATGGAGATTAGATCCTTCTTAACAGAAGGGCGCAGTTCGGCTGGATCTCG
34_B            ACTTCTTGACATGGAGATTAGATCCTTCTTAACAGAAGGGCGCAGTTCGGCTGGATCTCG
45_B            ACTTCTTGACATGGAGATTAGATCCTTCTTAACAGAAGGGCGCAGTTCGGCTGGATCTCG
26_D            ACTTCTTGACATGGAGATTAGATCCTTCTTAACAGAAGGGCGCAGTTCGGCTGGATCTCG
35_D            ACTTCTTGACATGGAGATTAGATCCTTCTTAACAGAAGGGCGCAGTTCGGCTGGATCTCG
                ************************************************************

3A_M__C         CACAGGTGCTGCATGGCTGTCGTCAGCTCGTGTCGTGAGATGTTGGGTTAAGTCCCGCAA
8_E             CACAGGTGCTGCATGGCTGTCGTCAGCTCGTGTCGTGAGATGTTGGGTTAAGTCCCGCAA
2_A             CACAGGTGCTGCATGGCTGTCGTCAGCTCGTGTCGTGAGATGTTGGGTTAAGTCCCGCAA
43_B            CACAGGTGCTGCATGGCTGTCGTCAGCTCGTGTCGTGAGATGTTGGGTTAAGTCCCGCAA
34_B            CACAGGTGCTGCATGGCTGTCGTCAGCTCGTGTCGTGAGATGTTGGGTTAAGTCCCGCAA
45_B            CACAGGTGCTGCATGGCTGTCGTCAGCTCGTGTCGTGAGATGTTGGGTTAAGTCCCGCAA
26_D            CACAGGTGCTGCATGGCTGTCGTCAGCTCGTGTCGTGAGATGTTGGGTTAAGTCCCGCAA
35_D            CACAGGTGCTGCATGGCTGTCGTCAGCTCGTGTCGTGAGATGTTGGGTTAAGTCCCGCAA
                ************************************************************

3A_M__C         CGAGCGTAACCCTCATCCTTAGTTGCCAGCGGGTTAAGCCGGGCACTTTAAGGAGACTGC
8_E             CGAGCGTAACCCTCATCCTTAGTTGCCAGCGGGTTAAGCCGGGCACTTTAAGGAGACTGC
2_A             CGAGCGTAACCCTCATCCTTAGTTGCCAGCGGGTTAAGCCGGGCACTTTAAGGAGACTGC
43_B            CGAGCGTAACCCTCATCCTTAGTTGCCAGCGGGTTAAGCCGGGCACTTTAAGGAGACTGC
34_B            CGAGCGTAACCCTCATCCTTAGTTGCCAGCGGGTTAAGCCGGGCACTTTAAGGAGACTGC
45_B            CGAGCGTAACCCTCATCCTTAGTTGCCAGCGGGTTAAGCCGGGCACTTTAAGGAGACTGC
26_D            CGAGCGTAACCCTCATCCTTAGTTGCCAGCGGGTTAAGCCGGGCACTTTAAGGAGACTGC
35_D            CGAGCGTAACCCTCATCCTTAGTTGCCAGCGGGTTAAGCCGGGCACTTTAAGGAGACTGC
                ************************************************************

3A_M__C         CAGTGGTAAACTGGAGGAAGGTGGGGATGATGTCAAGTCAGCACGGCCCTTATGGGGTGG
8_E             CAGTGGTAAACTGGAGGAAGGTGGGGATGATGTCAAGTCAGCACGGCCCTTATGGGGTGG
2_A             CAGTGGTAAACTGGAGGAAGGTGGGGATGATGTCAAGTCAGCACGGCCCTTATGGGGTGG
43_B            CAGTGGTAAACTGGAGGAAGGTGGGGATGATGTCAAGTCAGCACGGCCCTTATGGGGTGG
34_B            CAGTGGTAAACTGGAGGAAGGTGGGGATGATGTCAAGTCAGCACGGCCCTTATGGGGTGG
45_B            CAGTGGTAAACTGGAGGAAGGTGGGGATGATGTCAAGTCAGCACGGCCCTTATGGGGTGG
26_D            CAGTGGTAAACTGGAGGAAGGTGGGGATGATGTCAAGTCAGCACGGCCCTTATGGGGTGG
35_D            CAGTGGTAAACTGGAGGAAGGTGGGGATGATGTCAAGTCAGCACGGCCCTTATGGGGTGG
                ************************************************************

3A_M__C         GCTACACACGTGCTACAATGGTGACTACAATAGGTTGCAATGTCGTAAGGCTGAGCTAAT
8_E             GCTACACACGTGCTACAATGGTGACTACAATAGGTTGCAATGTCGTAAGGCTGAGCTAAT
2_A             GCTACACACGTGCTACAATGGTGACTACAATAGGTTGCAATGTCGTAAGGCTGAGCTAAT
43_B            GCTACACACGTGCTACAATGGTGACTACAATAGGTTGCAATGTCGTAAGGCTGAGCTAAT
34_B            GCTACACACGTGCTACAATGGTGACTACAATAGGTTGCAATGTCGTAAGGCTGAGCTAAT
45_B            GCTACACACGTGCTACAATGGTGACTACAATAGGTTGCAATGTCGTAAGGCTGAGCTAAT
26_D            GCTACACACGTGCTACAATGGTGACTACAATAGGTTGCAATGTCGTAAGGCTGAGCTAAT
35_D            GCTACACACGTGCTACAATGGTGACTACAATAGGTTGCAATGTCGTAAGGCTGAGCTAAT
                ************************************************************

3A_M__C         CCGTAAAAGTCATCTCAGTTCGGATTGTCCTCTGCAACTCGAGGGCATGAAGTCGGAATC
8_E             CCGTAAAAGTCATCTCAGTTCGGATTGTCCTCTGCAACTCGAGGGCATGAAGTCGGAATC
2_A             CCGTAAAAGTCATCTCAGTTCGGATTGTCCTCTGCAACTCGAGGGCATGAAGTCGGAATC
43_B            CCGTAAAAGTCATCTCAGTTCGGATTGTCCTCTGCAACTCGAGGGCATGAAGTCGGAATC
34_B            CCGTAAAAGTCATCTCAGTTCGGATTGTCCTCTGCAACTCGAGGGCATGAAGTCGGAATC
45_B            CCGTAAAAGTCATCTCAGTTCGGATTGTCCTCTGCAACTCGAGGGCATGAAGTCGGAATC
26_D            CCGTAAAAGTCATCTCAGTTCGGATTGTCCTCTGCAACTCGAGGGCATGAAGTCGGAATC
35_D            CCGTAAAAGTCATCTCAGTTCGGATTGTCCTCTGCAACTCGAGGGCATGAAGTCGGAATC
                ************************************************************

3A_M__C         GCTAGTAATCGTGGATCAGCATGCCACGGTGAATACGTTCTCGGGTCTTGTACACACTGC
8_E             GCTAGTAATCGTGGATCAGCATGCCACGGTGAATACGTTCTCGGGTCTTGTACACACTGC
2_A             GCTAGTAATCGTGGATCAGCATGCCACGGTGAATACGTTCTCGGGTCTTGTACACACTGC
43_B            GCTAGTAATCGTGGATCAGCATGCCACGGTGAATACGTTCTCGGGTCTTGTACACACTGC
34_B            GCTAGTAATCGTGGATCAGCATGCCACGGTGAATACGTTCTCGGGTCTTGTACACACTGC
45_B            GCTAGTAATCGTGGATCAGCATGCCACGGTGAATACGTTCTCGGGTCTTGTACACACTGC
26_D            GCTAGTAATCGTGGATCAGCATGCCACGGTGAATACGTTCTCGGGTCTTGTACACACTGC
35_D            GCTAGTAATCGTGGATCAGCATGCCACGGTGAATACGTTCTCGGGTCTTGTACACACTGC
                ************************************************************

3A_M__C         CCGTCACGCCATGGGAATTGGCTTAACTCGAAGCTGGTGCGCCAACCGCAAGGAGGCAGC
8_E             CCGTCACGCCATGGGAATTGGCTTAACTCGAAGCTGGTGCGCCAACCGCAAGGAGGCAGC
2_A             CCGTCACGCCATGGGAATTGGCTTAACTCGAAGCTGGTGCGCCAACCGCAAGGAGGCAGC
43_B            CCGTCACGCCATGGGAATTGGCTTAACTCGAAGCTGGTGCGCCAACCGCAAGGAGGCAGC
34_B            CCGTCACGCCATGGGAATTGGCTTAACTCGAAGCTGGTGCGCCAACCGCAAGGAGGCAGC
45_B            CCGTCACGCCATGGGAATTGGCTTAACTCGAAGCTGGTGCGCCAACCGCAAGGAGGCAGC
26_D            CCGTCACGCCAT------------------------------------------------
35_D            CCGTCACGCCATGGGAATTGGCTTAACTCGAAGCTGGTGCGCCAACCGCAAGGAGGCAGC
                ************

3A_M__C         CATTTAAGGTTGGGTCAGTGACTAGGTGA
8_E             CATTTAAGGTTGGGTCAGTGACTAGGTGA
2_A             CATTTAAGGTTGGGTCAGTGACTAGGGTG
43_B            CATTTAAGGTTGGGTCAGTGACTAGG---
34_B            CATTTAAGGTTGGGTCAGTGACTAGGGTG
45_B            CATTTAAGGTTGGGTCAGTGACTAGGGTG
26_D            -----------------------------
35_D            CATTTAAGGTTGGGTCAGTGACTAGGGTG
```

**S1 Fig: Multiple sequence alignment of eight full-length *Anaplasma* 16S rRNA genes.** These sequences from test mouse and seven naturally infected camels were 100% identical and BLAST of GenBank showed highest identity of 100% to “*Candidatus* Anaplasma camelii” (MUSCLE v3.8). 3A_M_C = *“Ca.* Anaplasma camelii*”* sequenced in test mouse; 8E - 35D = *“Ca.* Anaplasma camelii*”* detected in camels.
